# SPIRE, Surface Projection Image Recognition Environment for bicontinuous phases: application for plastid cubic membranes

**DOI:** 10.1101/2021.04.28.441812

**Authors:** Tobias M. Hain, Michał K. Bykowski, Matthias Saba, Myfanwy E. Evans, Gerd E. Schröder-Turk, Łucja M. Kowalewska

**Author notes:** ^*^For correspondence (GEST); (ŁK).

## Abstract

Bicontinuous membranes in cell organelles epitomise nature’s ability to create complex functional nanostructures. Like their synthetic counterparts, these membranes are characterised by continuous membrane sheets draped onto topologically complex saddle-shaped surfaces with a periodic network-like structure. In cell organelles, their structure sizes around 50–500 nm and fluid nature make Transmission Electron Microscopy (TEM) the analysis method of choice to decipher nanostructural features. Here we present a tool to identify bicontinuous structures from TEM sections by comparison to mathematical “nodal surface” models, including the hexagonal lonsdaleite geometry. Our approach, following pioneering work by Deng and Mieczkowski (1998), creates synthetic TEM images of known bicontinuous geometries for interactive structure identification. We apply the method to the inner membrane network in plant cell chloroplast precursors and achieve a robust identification of the bicontinuous diamond surface as the dominant geometry in several plant species. This represents an important step in understanding their as yet elusive structure-function relationship.

## Introduction

Biological membranes, dynamic yet stable unique assemblies of lipids and proteins, are selective barriers and enzymatically active regions playing a crucial role in orchestrated cells’ functioning. From a structural point of view, they are described mainly as flat sheets or small folded isolated entities called vesicles. Interestingly, in specific cases, almost all types of cellular membranes can form symmetrical, bicontinuous configurations called “cubic membranes” (***Almsherqi et al., 2006, 2009***). These are characterised by a spatial structure based on a periodic network- or labyrinth-like geometry, defined by uninterrupted negatively-curved membranes and with high symmetry (often cubic) (***Bouligand, 1990***; ***Luzzati, 1997***). These can be modelled by negatively curved surfaces as the spatial model for the bilayer membrane. Note, that herein the phrase “cubic membrane” is used synonymously for any bicountinous membranes with two membrane-separated aqueous channels and with a cubic or otherwise highly-symmetric spatial structure.

Cubic membranes are observed in cells of different organisms, from protozoa to mammals. They can self-organize from almost all types of membranes, including, e.g., endoplasmic reticulum (ER), plasma membrane, mitochondria and plastid inner membranes, and inner nuclear membrane (reviewed in ***Almsherqi et al., 2009***). Due to the length-scale of such structures with typical periodicities between 50–500 nm our knowledge about cubic membrane arrangements is almost exclusively obtained from electron microscopy data. Note that this is in contrast to bicontinuous soft matter phases, with much smaller periodicity, where X-ray and neutron scattering has been traditionally used to identify structures.

Although the highly regular nature of the membrane arrangements has been noted by many authors, structure identification remains difficult, due to a lack of widely available image processing tools for this purpose. As a result many of these structures have been inaccurately identified as, for example, tubular inclusions, undulating membranes, or cisternal systems (for the review of this issue see ***Almsherqi et al., 2006, 2009***; ***Cui et al., 2020***), instead of associating them correctly with cubic phases. Already in the 1960s researchers recognized the possibility of non-lamellar phase formation by cellular membranes (e.g. ***Cheville, 1966***; ***Eymé, 1963***; ***Folliot and Maillet, 1965***; ***Lang and Rae, 1967***; ***Pappas and Brandt, 1959***; ***Schuster, 1965***; ***Wooding, 1967***). However, the deep understanding of the factors governing the formation of such complex amphiphilic arrangements is still unknown, partially due to the scattered nature and incorrect annotations of reported data. On the other hand, considerable advancement in recognition of intrinsic and extrinsic factors playing a role in elementary membrane bending and curvature sensing has been made, forming a solid foundation for further studies in understanding how the complex architecture of cubic membranes controls cellular traffic (***Assoian et al., 2019***; ***Callens et al., 2020***; ***Campelo et al., 2014***; ***Jarsch et al., 2016***; ***Kozlov et al., 2014***; ***Lou et al., 2018***; ***Pezeshkian and Marrink, 2021***; ***Raven, 2021***; ***Simunovic et al., 2019***; ***van Zanten and Mayor, 2015***; ***Zimmerberg and Kozlov, 2006***)

Recently there is a growing interest in the possibility of obtaining nature-inspired and, therefore, stable, large length-scaled cubic systems (>50 nm) to develop concepts to tackle different multidisciplinary issues and healthcare problems (reviewed in ***Mezzenga et al., 2019***). Naturally occurring cubic systems are an important inspiration for designing biomimetic nanostructures used in, e.g., drug delivery systems (***Mulet et al., 2013***; ***Porras-Gomez and Leal, 2019***), controlled release of solubilized molecules (***Clulow et al., 2020***), material science (***Han and Che, 2018***; ***Kresge et al., 1992***), and in the determination of protein structure (***Speziale et al., 2016***). However, such broad interdisciplinary interest in bicontinuous systems has not yet been addressed by recent fundamental research on naturally occurring cubic membranes. Key factors constraining advances in this field are the recognition and spatial analysis of observed membrane arrangements. Such data are crucial to establishing a model system for further biochemical studies and, finally, to discover the shape-dependent role of cubic membranes. This generates a great demand for methods to recognize and measure three-dimensional (3D) properties of periodic assemblies.

In terms of topology and geometry, cubic membrane structures can be described using triply-periodic minimal surfaces (TPMS). Minimal surfaces are surfaces that locally minimize their surface area based on some global constraint (e.g., a surface between a given boundary). As a consequence of this minimisation they have a mean curvature of zero at all points on the surface. Triply-periodic minimal surfaces are minimal surfaces with a crystalline, symmetric structure where they repeat in three independent translation directions in space. A wide array of triply-periodic minimal surfaces have been described mathematically with different crystallographic symmetry. However, three particular TPMS with cubic symmetry are most commonly observed in biological cubic membranes. Primitive and diamond types were characterized by Schwarz in 1865 (***Schwarz, 1890***) and the so-called gyroid recognized by ***Schoen*** (***1970***) almost a hundred years later (1970) (reviewed in ***Hyde et al., 1996***). TPMSs divide inner space into two separated, intertwining yet open channels. In terms of cubic membranes, the presence of such isolated regions of given sizes might have tremendous consequences in constraining molecular motion. Moreover, cubic membranes and other highly symmetric structures, such as the Lonsdaleite or Schwarz’ Hexagonal surface, generally based on defined TPMS templates, can form multilamellar systems leading to the formation of additional channels of potentially different composition and, therefore, function (e.g. ***Almsherqi et al., 2012***).

One of the examples of extensively studied cubic membrane assemblies is a prolamellar body (PLB) of plant etioplasts, see ***Figure 1***. The PLB is a direct precursor of the chloroplast thylakoid network, and their lipid-pigment-protein composition shares some similarities (for the review of PLB composition, see: ***Adam et al., 2011***; ***Kowalewska et al., 2019***; ***Pribil et al., 2014***). The PLB is considered as a lipid reservoir during tubular-lamellar transition increasing the efficiency of grana formation (***Armarego-Marriott et al., 2019***; ***Pipitone et al., 2021***). The number of PLB building blocks play a crucial role in maintaining its structure, including protochlorophyllide:light-dependent protochlorophyllide oxidoreductase:NADPH complex as well as particular galactolipids and carotenoids (***Floris and Kühlbrandt, 2021***; ***Bykowski et al., 2020***; ***Cazzonelli et al., 2020***; ***Franck et al., 2000***; ***Fujii et al., 2019***; ***Nguyen et al., 2021***; ***Sperling et al., 1998***). Although the role of light-dependent protochlorophyllide oxidoreductase in membrane tubulation was proven recently using electron cryo-tomography techniques (***Nguyen et al., 2021***; ***Floris and Kühlbrandt, 2021***), factors governing the transition of tubular arrangements into cubic configuration remain elusive (***Wietrzynski and Engel, 2021***). The role of already recognized structure-remodeling proteins of plastid membranes (CURT1, VIPP1) in this process cannot be excluded and require further investigation (***Armbruster et al., 2013***; ***Gupta et al., 2020***). PLBs are rare examples of cubic membranes resembling an imbalanced structure, in which the two aqueous channels are geometrically different; the smaller channel is a direct precursor of thylakoid lumen of chloroplasts (***Kowalewska et al., 2016***). The PLB structural organization’s variability has been studied extensively revealing different possible geometrical forms of this intricate periodic pattern. In early studies, PLB structures were most frequently referred to zinc sulfide crystal forms of wurtzite (lonsdaleite) and zincblende type, both based on tetrahedral units forming complex hexagonal networks and as such with hexagonal instead of cubic symmetry. The primitive, face-centered diamond and double diamond cubic membrane types were also proposed (***Gunning, 1965***; ***Gunning and Steer, 1975***; ***Ikeda, 1968***; ***Landh, 1996***; ***Menke, 1963***). These variable structural annotations were made based on the analyses of randomly cutted PLB sections visible in two-dimensional (2D) TEM micrographs via their comparison with 3D models (physical or rendered) of mentioned structures. However, even comparing many 2D projections of a 3D structure (TEM specimen) at different viewing angles with an actual 3D model is not directly verifiable and probably led to such inconsistency in the identification of the most abundant spatial PLB configuration.

**Figure 1.**
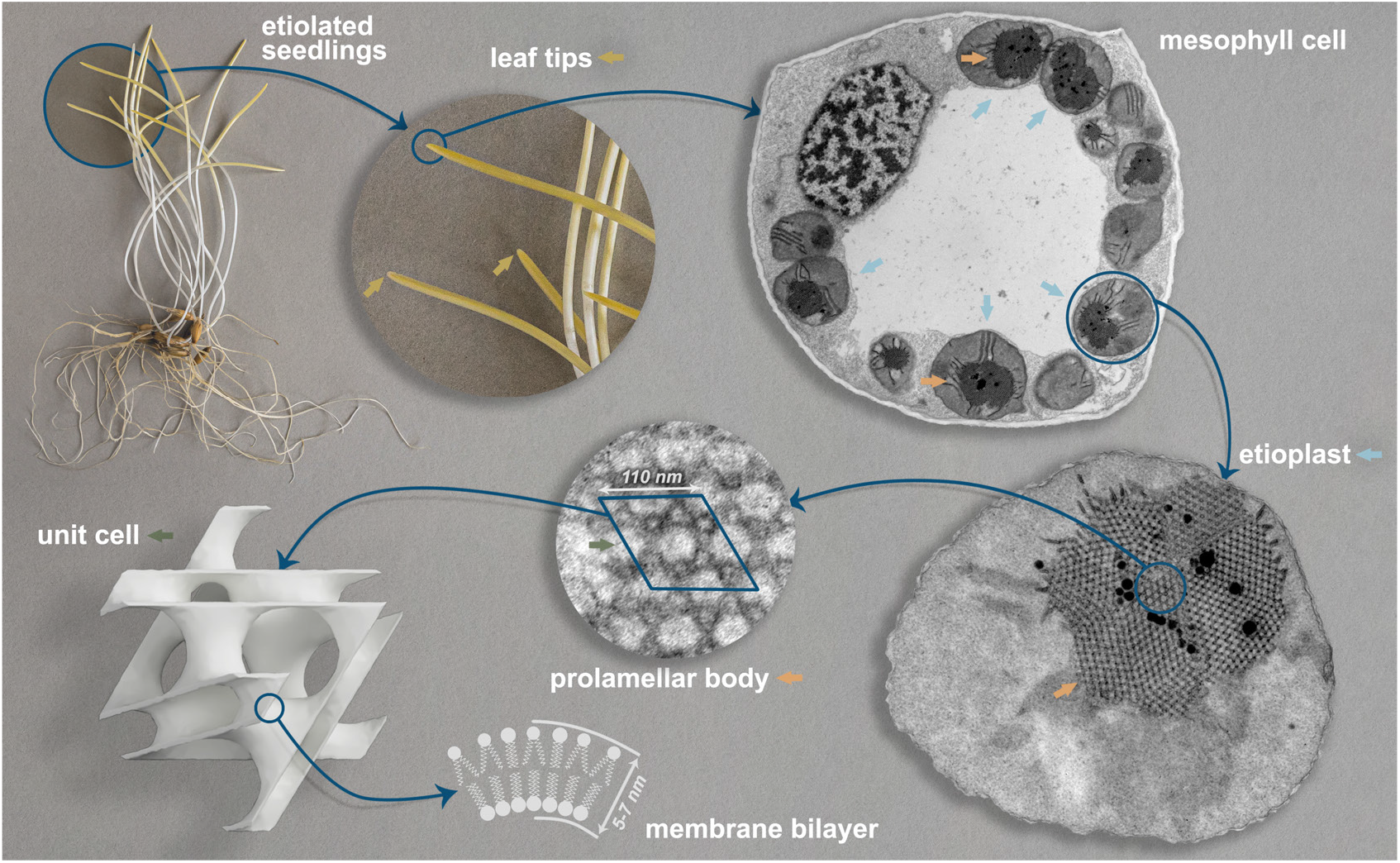
Cellular localization of cubic membrane assembly – prolamellar body (PLB) in etiolated seedlings of oat *Avena sativa*. Cubic structure of PLBs located in etiolated seedlings of angiosperms – here exemplified by *Avena sativa* growing for 7 days in complete darkness. PLBs develop in etioplasts present in developing leaves’ mesophyll cells (yellow parts of the seedling visible above white part of the shoot). The cubic arrangement of the PLB is characterized by an imbalanced proportion of water channels and a relatively small length-scale compared to other naturally-occurring cubic membranes; a single membrane separates both sides of the structure. Note that recent results indicated that PLB membrane is densely decorated with light-dependent protochlorophyllide oxidoreductase protein, which role in membrane tubulation has been proven in *in vitro* studies (***Nguyen et al., 2021***; ***Floris and Kühlbrandt, 2021***). Apart from marked elements, the presented photographs, and electron micrographs are not shown to scale; mesophyll cell and etioplast are free-form selected manually from Transmission Electron Microscopy (TEM) images of etiolated oat leaves.

There are two main methods for analyzing the 3D structure of PLB and other cubic membrane arrangements. Both methods are based on the assumption that cubic membrane structures correspond directly to mathematically well-defined TPMSs. The first approach consists of the visualization of cubic membranes using electron tomography, further segmentation, modeling the periodic arrangement, and finally, its direct comparison to the rendered 3D model of different bicontinuous structures with variable surface parameters and length-scales (***Chong and Deng, 2012***; ***Demurtas et al., 2015***; ***Kowalewska et al., 2016***). Such a method is time, money, and computational power-consuming due to the operation on the 3D objects. It is also limited to the cubic membrane structures of particular length scales. Cubic membranes of small periodicities cannot be reconstructed with sufficient resolution from the electron tomography data. Those with large periodicities (around 500 nm) exceed the typical electron tomography sample’s thickness. Moreover, it should be stressed that manual segmentation of cubic membranes is complicated, and auto-mated methods, while sufficient to estimate general structural parameters (e.g., channel volume or surface area), fail in terms of precise shape visualization.

Alternatively, the second method is performed using 2D TEM images of cubic membranes and their direct comparison with a simulation of a 2D TPMS projection of given parameters. The idea of the “template matching” method for cubic structure recognition was initially introduced by Mark Mieczkowski and Yuru Deng (***Deng and Mieczkowski, 1998***; ***Deng et al., 1999***). It was successfully implemented to recognize the surface type of several membrane arrangements, e.g., in mitochondria of starved *Chaos carolinensis* (***Deng et al., 1999***) or chloroplasts of green alga *Zygnema* sp. in the log phase of growth (***Zhan et al., 2017***) using the developed software called QMSP (Cubic Membrane Simulation Program). The QMSP tool enabled the generation of a library of projection images for structures of primitive, double diamond, and gyroid surfaces; the user could manipulate the direction of the projection, number of visualized unit cells (UCs), and thickness of the projected TPMS region. The tool is not publicly available and has limited functionalities in projection scaling, surface types, channel balance, and measurement properties.

Inspired by ***Deng and Mieczkowski*** (***1998***), we introduce SPIRE (Surface Projection Image Recognition Environment), a new open-source tool to simulate TEM images of TPMSs. It addresses afore-mentioned issues, vastly extends and improves the structure identification process by focusing on interactive matching and lay the foundation for an automated identification process.

The SPIRE tool focuses on the interactive matching of TEM images, with access to a large range of parameters and metrics of the structure. It is broadly applicable to reliably recognize and analyze structural, spatial properties of bicontinuous arrangements visualized in electron microscopy. A widespread representation of cubic membranes in living organisms highlights the importance of our tool for a large community of biologists. The intuitive SPIRE graphical user interface (GUI), see ***Figure 14***, can be used by a broad group of scientists, including those with no explicit knowledge of the geometrical description of cubic structures.

Although we developed SPIRE to investigate cellular cubic membranes visualized in TEM images, it is also a suitable tool for analyzing microscopy images of any highly symmetric arrangements such as cuboids or polymer assemblies. In the latter, we see similar geometries to those found in the biological cubic membranes (***Bates, 2005***; ***Kirkensgaard et al., 2011***; ***Han et al., 2020***) or cubic rod packings (***O’Keeffe et al., 2001***). The latter are used to model the keratin microstructure in skin cells, a geometry that is closely related to—and likely coexistent with—a gyroid surface (***Evans and Hyde, 2011***; ***Evans and Roth, 2014***; ***Norlén and Al-Amoudi, 2004***). Synthetic cubic structures are mainly analyzed using scattering methods, but in non-uniform samples, additional microscopy analyses are required. Moreover, SPIRE could be used to examine scanning electron microscopy (SEM) images of, e.g., non-membrane based naturally occurring highly symmetric arrangements such as gyroid recognized in wing scales of a butterfly – *Callophrys rubi* (***Saranathan et al., 2010***; ***Schröder-Turk et al., 2011***) or diamond in beetle scales - *Lamprocyphus augustus* (***Galusha et al., 2008***).

This article first describes the main concepts and algorithms of the software, followed by a detailed walkthrough of a structure identification process using the PLB arrangements in etiolated seedlings as an example. The Appendix contains a video tutorial presenting the tool and workflow, as well as information about the built-in structures and parameters used throughout this article.

## Results

### Numerical procedures

***Figure 2*** illustrates the major steps in the simulation process used in SPIRE. This software simulates TEM images of ultrathin sections of biological samples, which are essentially planar projections of the structure inside the sample. In general, a planar projection is a flat, thus 2D image of a 3D structure. Though this transformation to a lower dimensional space causes a loss of information, in many cases it yields an image providing a better overview than the 3D structure.

**Figure 2.**
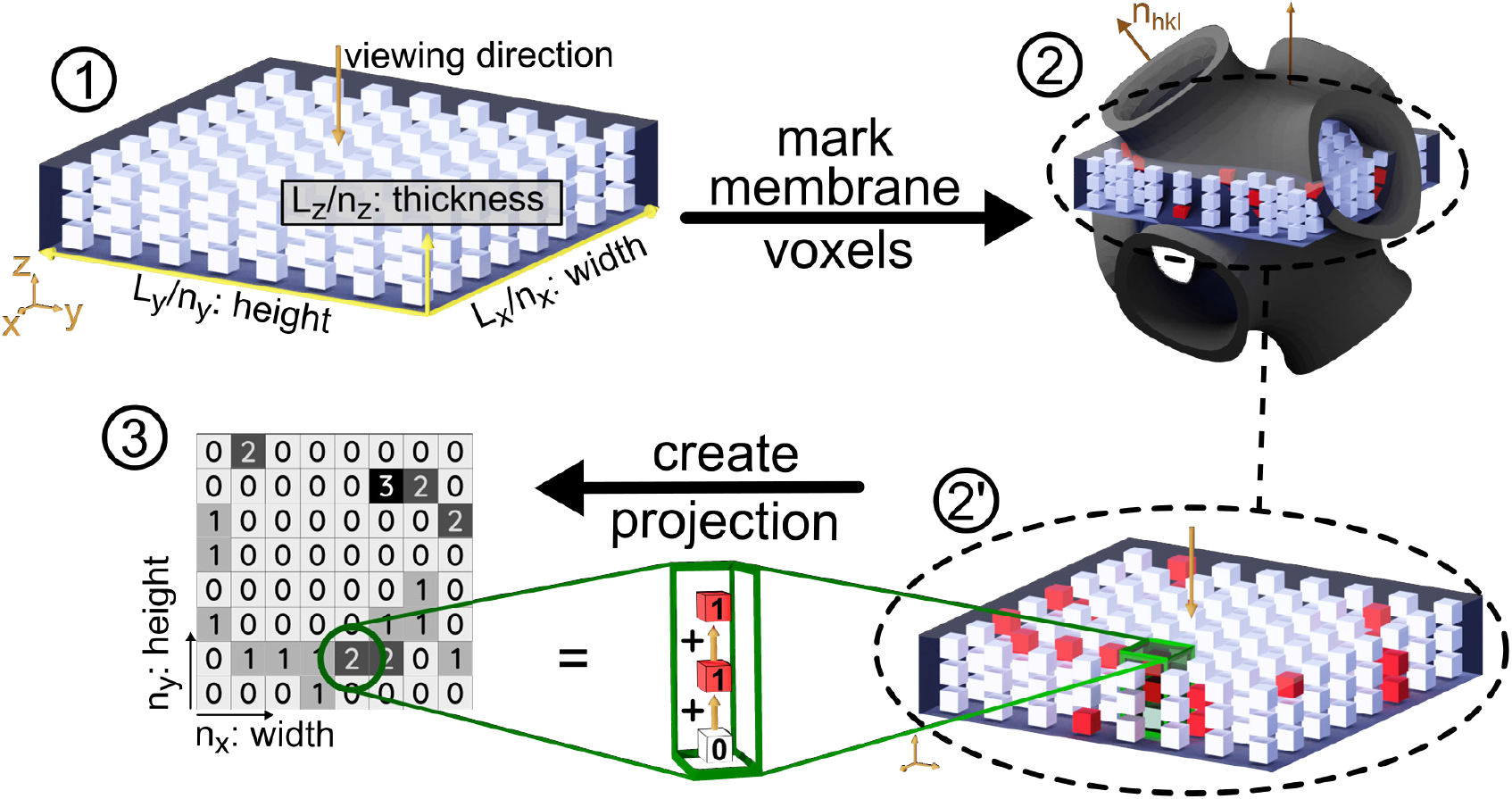
The basic steps in the process of computing a planar projection. (**1**) Schematic representation of the rectangular simulation box (called slice) with the dimensions (*L*_*x*_, *L*_*y*_, *L*_*z*_), filled with a grid of *n*_*x*_ · *n*_*y*_ · *n*_*z*_ voxels of size (*L*_*x*_/*n*_*x*_, *L*_*y*_/*n*_*y*_, *L*_*z*_/*n*_*z*_). The viewing direction is perpendicular to the *L*_*x*_*L*_*y*_ plane. Note that the space between voxels is just for visualisation and is not existent in the simulation. (**2**) All voxels are evaluated using the mathematical model of the surface structure: the slice is superimposed in the oriented membrane structure; a voxel is marked if located within a membrane (**2’**) The projection is computed by adding the values of voxels with identical and coordinates, i.e. all voxels congruent in viewing direction. Each marked voxel is valued as “1”, whereas unmarked voxels do not contribute (**3**) The simulated planar projection, a pixel image where each pixel brightness holds the number of marked, thus membrane, voxels. Its resolution is determined by the number of voxels (*n*_*x*_, *n*_*y*_) in the initial slice

Here, we aim for a projection emulating a TEM image of a 3D structure. The latter encodes the amount of beam attenuating material along its path that generates the microscope image: whereas dark regions in the image represent areas of a large amount of material along the beam path, brighter regions present less material.

To simulate those projections, a model of the sample is created by using one or multiple membranes, modeled as minimal or negatively curved surfaces. A discrete grid of points, where each point is either marked as “attenuating”, or not marked, that is “translucent”, is then used to compute the planar projection. In this section details on the underlying processes and methods will be presented, starting with the introduction of the mathematical model and description of the membrane structures used in the software.

### Mathematical modelling of bicontinous membranes

A mathematical model, the so-called nodal representation, is used to describe the membrane geometry. In this model, the true TPMSs are approximated by implicit functions 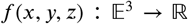, the so-called nodal functions (***von Schnering and Nesper, 1991***; ***Klinowski et al., 1996***). Surfaces are then defined using so-called level set parametrization: the surface is the set of all points where *f* (*x*, *y*, *z*) = *c*, where *c* is an arbitrary constant. Different values of *c* yield different surfaces. In case of the low genus TPMSs considered here, ***von Schnering and Nesper*** (***1991***);***Gandy et al.*** (***2001***) found that only a few terms in the nodal representation *f* (*x*, *y*, *z*) are sufficient for high accuracy with only little deviation from the exact minimal surface.

SPIRE contains nodal representations for several surfaces as well as rod packings. Three surfaces with cubic symmetries are included: the gyroid, diamond and primitive surface. The nodal representations of the latter are taken from ***von Schnering and Nesper*** (***1991***). We computed the nodal representation of the lonsdaleite (“hexagonal diamond”) surface from its corresponding spacegroup (SG194) and its structure factor (similar to ***Wohlgemuth et al., 2001***, and references therein). The leading term only representation reads

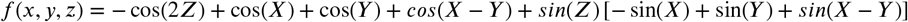

with

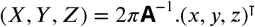

and **A** being the matrix comprising the three conical lattice vectors of a structure with hexagonal symmetry: 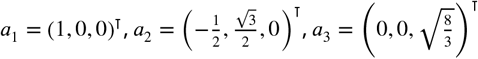 and the dot denoting the matrix product (see Crystallographic nature of highly symmetric membranes for more details). Note, that due to their repeating nature, all representations of triply-periodic surfaces can be expressed solely using periodic trigonometric functions. Although the presented formula yields a surface with a topology and geometry equivalent to the lonsdaleite structure, it is not a minimal surface. For the implementation in the software we therefore used a numerically optimised version^1^.

All TPMSs divide space into two intertwining channels. For this software, the nodal representation is chosen such as *f* (*x*, *y*, *z*) = 0 yields the balanced case: the membrane separates two channels of equal volume. Membranes at *f* (*x*, *y*, *z*) = *c*, where *c* is an arbitrary constant, separate two channels with unequal volumes. The constant *c* is thus a measure of the proportion of volumes of the two channels. The software allows to choose the position of the membrane either based on the constant *c* (“level set”) or the volume proportion of the two channels. Note, however, that only for membranes where *f* (*x*, *y*, *z*,) = 0 the nodal representation does approximate a true minimal surface^2^!

### Discretization of structure models

For the discretization of the structure models, an approach combining level-sets of the nodal representations with a distance-map is employed as follows. The process begins by computing the value of the nodal representation *f* (*p*_*x*_, *p*_*y*_, *p*_*z*_)for each voxel at position (*p*_*x*_, *p*_*y*_, *p*_*z*_) in the simulation box. As shown in ***Figure 2***, a voxel is marked, if *c*−*λ* < *f* (*p*_*x*_, *p*_*y*_, *p*_*z*_) < *c*+*λ*, where *λ* is a constant with the meaning of the width of the membrane. In a naive implementation, visually speaking a voxel is marked if it is located within the space bounded by the two level-set surfaces given by *f* (*x*, *y*, *z*) = *c* + *λ* and *f* (*x*, *y*, *z*) = *c* − *λ*. Note that due to the nature of the nodal representation, these two surfaces bounding the membrane are not necessarily parallel and thus would create a membrane with varying width. To resolve this issue, and being able to tune the width of the membrane in true length units, the following procedure is used: the parameter *λ* is chosen very small, such that ideally the resulting membrane width is only a single voxel thick. A so-called Euclidean distance map (*EDM*) (***Felzenszwalb and Huttenlocher, 2012***) is then computed. This function *EDM*(*p*_*x*_, *p*_*y*_, *p*_*z*_) assigns each voxel in the slice the value of the distance (in length units) of the current voxel to the closest marked one, in this case the closest membrane voxel. All voxels inside one channel are assigned a positive distance, whereas voxels in the other channel have negative distances to the membrane. The membrane with the desired width is then obtained by just marking all voxels where the value *EDM*(*p*_*x*_, *p*_*y*_, *p*_*z*_) is smaller than or equal to half of the membrane width.

Apart from these common single bilayer membrane structures, more complicated geometries such as double bilayer membranes occur in nature (***Deng and Mieczkowski, 1998***). To model such systems, the software allows multiple membranes to be added to the slice. Whereas the location of a single membrane is specified using the level-set value *c*, or the proportion of the channel volumes, the positions of additional membranes are given as a distance to the initial, level-set membrane. Additional parameters control the width of each membrane. Marking the voxels of additional membrane is conveniently done using the *EDM* computed before: for each membrane *i* of width _*i*_ and distance *d*_*i*_ all voxels for which *d*_*i*_ − (*w*_*i*_/2) < *EDM*(*p*_*x*_, *p*_*y*_, *p*_*z*_) < *d*_*i*_ + (*w*_*i*_/2) holds are marked.

Whereas a single bilayer membrane separates space into two channels, each additional membrane will add a further domain (called “channel” in the software; meaning the space between two parallel membranes). Since membranes have a finite width and thus a volume, this software internally considers the latter as channels, too. The total number of channels is then 2 · *n* + 1, where *n* is the number of membranes. The channels are labeled with increasing integers starting at “1” with the innermost channel (that contains the center of the unit cell). The innermost membrane will then have the channel number “2”, also view ***Figure 3*** for more information.

**Figure 3.**
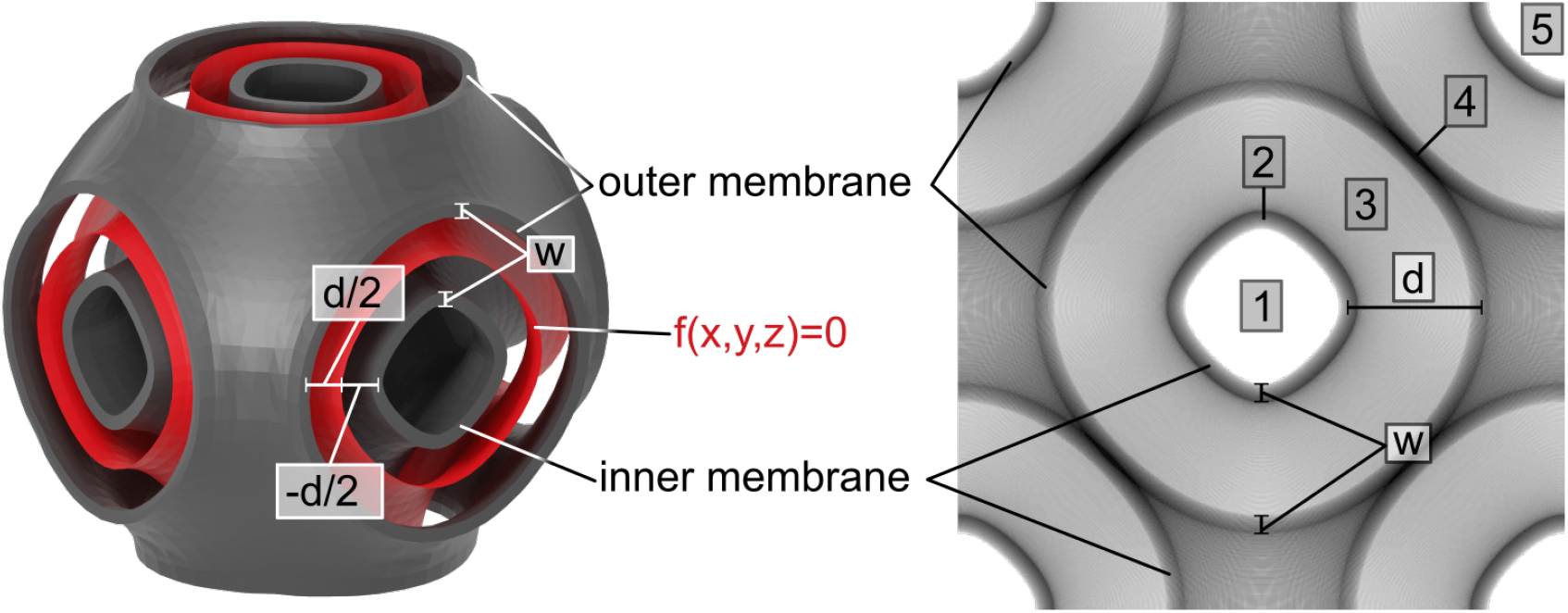
Multi-layer membrane structures and channel enumeration. A primitive surface multi-layer membrane system and its 2D projection in (100) orientation with two membranes of width *w* at a distance of *d* computed as parallel surfaces from the level-set membrane, the minimal surface at *f* (*x*, *y*, *z*) = 0, shown in red. The latter is only computed internally and does not show in the projection. The inner membrane is inside of the level-set membrane, therefore has a negative distance. The numbers denote the channel numbers of a total of 5 channels, of which 3 are “true” channels and 2 are membranes, also considered as channels internally.

So far, only systems related to the double “gyroid” (or other “double phases”) where a membrane (modelled as absorbing) separates two aqueous channels (modelled as translucent) were considered. These “double phases” are not the only structural models based on TPMSs. Several examples in nature, such as a gyroid surface in the wing scales of the butterfly *C. rubi* (***Schröder-Turk et al., 2011***) or a diamond surface in the beetle *L. augustus* (***Wilts et al., 2012***), have been reported with a single surface separating two intertwining channels, with one of the two filled by a solid material. To model such systems, the software allows for each individual channel to be filled with material. Then all voxels inside to the chosen channel are marked, i.e. modeled as opaque. Analogously, the software allows for membranes to be marked translucent to allow for more complex models or analyses.

### Crystallographic nature of highly symmetric membranes

After introducing the mathematical description of the TPMSs and their geometries, this description is now used to create synthetic images corresponding to the membrane structure in voxelized (3D pixel) form.

The simulation process starts with the initialization of the simulation box, a cuboid with dimensions *L*_*x*_, *L*_*y*_ and *L*_*z*_, where each edge is aligned with its corresponding axes of the canonical base, see ***Figure 2***. The simulation box will be called “slice” and represents the region of interest (*L*_*x*_ and *L*_*y*_) as well as physical thickness (*L*_*z*_) of the ultrathin section of tissue captured by the TEM image, and will be filled with a discretized version of the membrane structure. To store the latter, the simulation box is fitted with a regular, rectangular grid of a total of *n*_*x*_ · *n*_*y*_ · *n*_*z*_ sites, see part 1 in ***Figure 2***. On each site a small cuboid with dimensions (*dx*, *dy*, *dz*), a so-called voxel, is placed, such as there is no overlap or space between two adjacent voxels. Each of these voxels can be “unmarked”, i.e. translucent, or “marked”, i.e. opaque. In ***Figure 2*** this property is represented by the color “white” and “red” and will be assigned in a later step.

The slice dimension (*L*_*x*_, *L*_*y*_, *L*_*z*_) can be chosen freely by the user and is an important parameter to match the simulated projection to the size of the TEM image. That is, the slice dimensions should be chosen to correspond to the size of the TEM image of the section considered. The number of voxels in the slice can be tuned by providing the number of voxels in *x* (*n*_*x*_) and *z* (*n*_*z*_) direction. To obtain voxels with a square footprint (*dx* ≈ *dy*) the number of voxels in direction is then computed to *n*_*y*_ = *L*_*x*_/*L*_*y*_ · *nx* (or nearest integer). The voxel dimensions are computed automatically. The number of voxels determines the resolution and hence the quality of the projection, in line with the resolution of the TEM image.

The current version of the software is designed to model to periodic membrane structures. The latter are characterised by a translational unit cell (UC) of finite size and three vectors representing the periodic replication directions. That is, an infinitely large, continuous structure can be constructed by repeating the translational UC along the replication directions. Due to this repeating nature, points with identical geometry but different spatial coordinates, here called lattice points, can be identified. The black dots in ***Figure 4*** represent a possible choice of lattice points. For the structures considered in the tool, these lattice points represent locations on which the UC has to be positioned to recreate the infinite, continuous structure.

**Figure 4.**
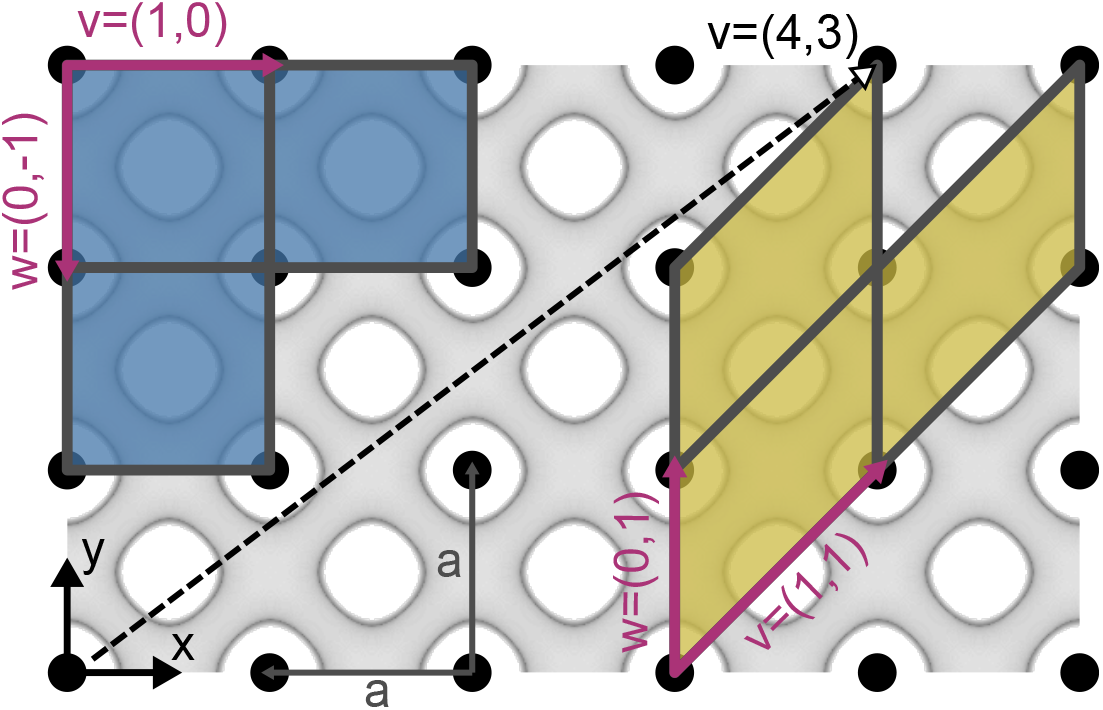
The definition of a unit cell (UC) and its base vectors. An example of a periodic structure in two-dimensional (2D) with two choices of lattice vectors and resulting UCs: the fundamental UC (smallest UC with least amount of information to fully replicate the infinite structure while being cubic or rectangular) and an inclination UC (see main text for definition). The black dots are a choice of lattice points and represent geometrically identical, since repeating locations in the structure. The choice of lattice vectors and thus UCs is arbitrary and the size of the UC depends on the choice of lattice vectors: for **v** = (1, 0) and **w** = (0, −1) the UC is square with an edge length of a (lattice constant), where as for **v** = (1, 1) and **w** = (0, 1) the UC is a parallelogram with edge lengths 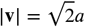 and |**w**| = *a*. The dashed vector *v* demonstrates that lattice vectors (and thus the UC) can get very long for odd directions.

The entire information of a periodic structure is thus stored in only a single translational UC and the three replication directions, called lattice vectors **u**, **v** and **w**. We conveniently choose these vectors as the bounding edges of the UC (see ***Figure 4***), although other choices exist and may be more fitting for different purposes. As a consequence the size and shape of the latter is solely determined by a choice of the three lattice vectors **u**, **v** and **w**, i.e. the replication directions.

Given a periodic structure, there is no unique choice of a UC: apart from the aforementioned constraint, the lattice vectors and thus the UC can be chosen arbitrarily. We introduce a distinct choice of the UC, the herein called fundamental UC. The latter is the cubic (for the surfaces with cubic symmetries, e.g., the primitive, diamond and gyroid surface) or rectangular (for rectangular symmetries, as the Lonsdaleite) UC which among all choices has the smallest volume and content possible (in ***Figure 4*** exactly one lattice point) while still containing all information needed to reproduce the entire structure. In the case of cubic or rectangular TPMSs the lattice vectors **u**, **v** and **w**, with |**u**| = *c*, |**v**| = *a* and |**w** = *b*, of the fundamental UC can be conveniently aligned with the Cartesian coordinate axes **x**, **y** and **z**. Any UC with a different shape or choice of lattice vectors, especially rotated versions of the fundamental UC, will herein be called inclination UC. The appendix lists all fundamental unit cells of the structures built into the software in section Fundamental unit cells of built-in surfaces.

Whereas in real, biological samples the orientations of the TPMS structures are in most cases random^3^, the orientation of the membrane structure in the slice can be fixed in the simulation, providing the ability to generate projections with identified orientations. The fixed viewing direction (electron/light beam) onto the sample in TEM is reflected in the software by fixing the viewing direction on to the 3D slice arbitrarily but conveniently to the negative z-direction, i.e. perpendicular to the *L*_*x*_*L*_*y*_ plane of the slice, as shown in panel 1 in ***Figure 2***. This choice will facilitate the subsequent step of computing the planar projection.

The orientation of the membrane structure within the slice is described by using the so-called Miller indices (*hkl*) (***Kittel, 2004***), a triplet of integer numbers denoting the orientation of a lattice plane. The Miller indices can be defined by *h* = *p/S*_*x*_, *k* = *p/S*_*y*_ and *l* = *p/S*_*z*_, where *p* is the smallest integer, for which (*hkl*) is a triplet of integers without a common divisor and *S*_*i*_ is the point where the plane intersects the *i*-th UC base vector (equivalent to coordinate axes in cubic symmetries), given in multiples of the respective lattice vector, see ***Figure 5***. Thus the Miller indices define a plane fixed by three points in space. An index value of 0 means the plane is parallel to the respective axis. For structures with a rectangular fundamental UC, that is all of the implemented surfaces, a more intuitive approach to the Miller indices is to interpret them as components of the normal vector **n**_*hkl*_ on the plane they are denoting: **n**_*hkl*_ = ((1/*a*) · *h*, (1/*b*) · *k*, (1/*c*) · *l*)^⊺^. Note that we use a rectangular fundamental UC for cubic structures and the hexagonal Lonsdaleite alike. The latter choice implies that the (*hkl*) values do not correspond to the expected direction in the crystallographic convention and extra care needs to be taken when specifying the inclination. The appendix Fundamental unit cells of built-in surfaces lists the exact dimensions and choice of lattice vectors of all fundamental unit cells in the software. Neglecting scaling factors, since only the orientation of the normal vector is of relevance, yields for a cubic UC (as is the case for the primitive, diamond and gyroid surface) **n**_*hkl*_ = (*h, k, l*)^⊺^.

**Figure 5.**
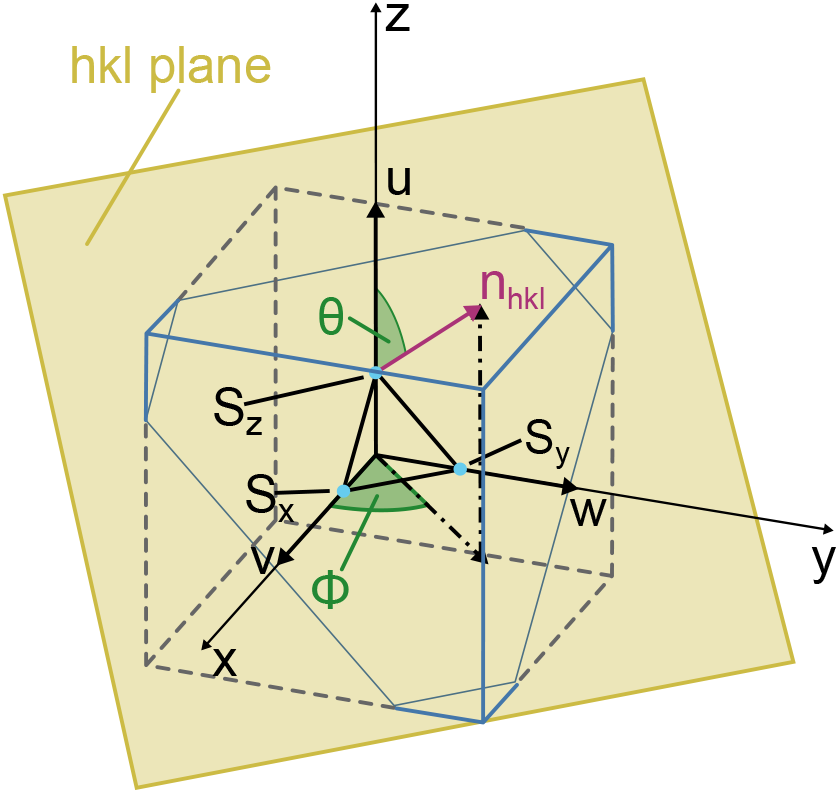
Definition of Miller indices and orientation denotation. *S*_*x*_, *S*_*y*_, *S*_*x*_ are the points, given in terms of the lattice vectors **v**, **w**, **u**, where the plane, denoted by the Miller indices (*hkl*), intersects the **x**,**y**,**z** axes (***Kittel, 2004***). For structures with cubic or rectangular fundamental unit cells (i.e. all structures implemented in the tool), the normal vector **n**_*hkl*_ on the (*hkl*) plane—and thus a direction—can be written in terms of the Miller indices: **n**_*hkl*_ = ((1/*a*) · *h,* (1/*b*) · *k,* (1/*c*) · *l*)^⊺^. Formally *h*,*k*,*l* are defined by *h* = *p/S*_*x*_, *k* = *p/S*_*y*_ and *l* = *p/S*_*z*_ where *p* is the smallest integer, for which (*hkl*) is a triplet of integers without a common divisor. The polar angle Φ is the angle between the **x** axis and the projection of **n**_*hkl*_ onto the -plane, the azimuthal angle e is the angle between the **z** axis and **n**_*hkl*_.

Internally the orientation of the normal vector given as Miller indices is converted in two angles, the polar angle Φ and the azimuthal angle Θ, where Φ denotes the angle between the **x** axis and the projection of **n**_*hkl*_ onto the *xy*-plane and Θ the angle between the **z** axis and **n**_*hkl*_. See ***Figure 5*** for a graphical representation.

In SPIRE the orientation of the fundamental UC, where **v**, **w**, **u** align with the coordinate axes, is assigned the “neutral” orientation (001) with **n**_001_=**z**, i.e. the orientational normal vector **n**_*hkl*_ aligns with the positive **z** direction. The two orientation angles then are Φ = 0 and Θ = 0.

In order to orient the structure inside the slice, a triplet of Miller indices is provided by the user, defining the viewing angle on the structure as vector **n**_*hkl*_, as discussed above. However, since the viewing angle in the tool is fixed to the **z** axis, the vector **n**_*hkl*_ needs to be aligned with the **z** axis, applying the same rotations on the membrane structure, as visualized in ***Figure 6***.

**Figure 6.**
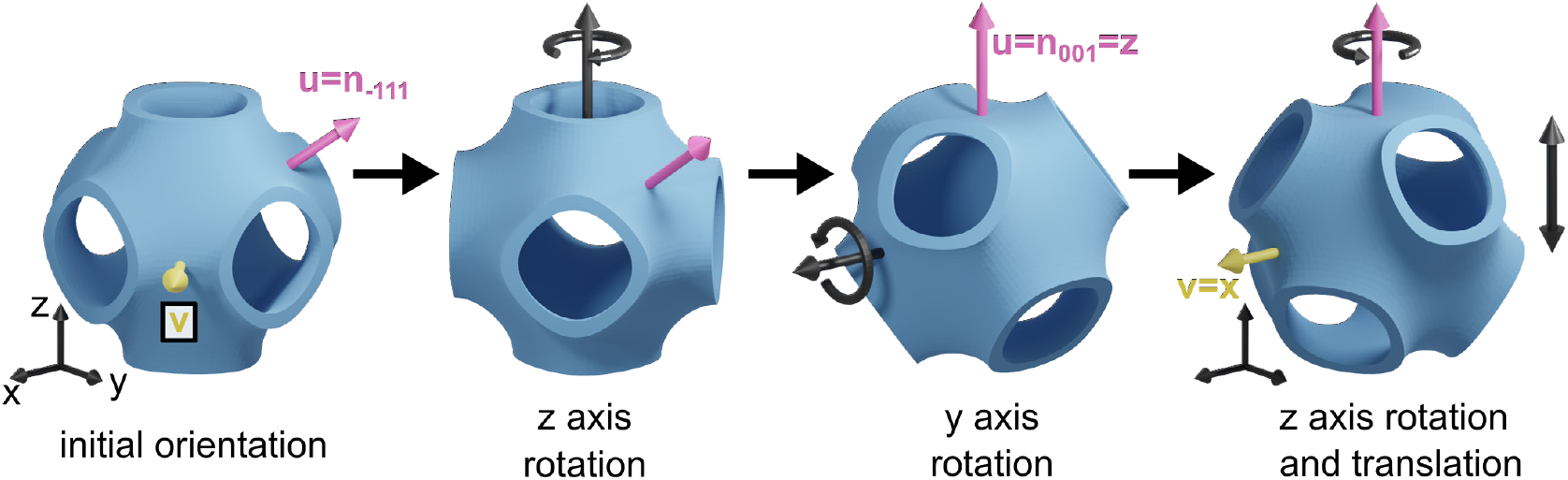
Orienting the membrane structure in the virtual sample. To simulate a projection with the desired viewing angle, the structure needs to be inside the slice. Starting at the orientation of the fundamental unit cell (UC), the structure is first rotated around the **z** axis by the polar angle Φ, rotating the vector **u** into the -plane, and subsequently around the **y** axis by the azimuthal angle Θ, aligning **u** with the **z** axis and thus the viewing direction. The last rotation around the **z** axis aligns the in-plane vector **v** with the **x** axis, such that the inclination UC has minimal volume given the normal vector **n**_*hkl*_. In a last step the structure can be translated along the **z** axis to choose the termination.

The alignment is performed in three separate rotations and a final translation, as shown in ***Figure 6***. The TPMS structure is initialized as the fundamental UC, with the lattice vector **u** aligned with **n**_*hkl*_. The TPMS structure is then rotated around the **z** axis by the angle Φ, and subsequently around the **y** axis by the angle Φ. Now the lattice vector **u** is aligned with the **z** axis. i.e. the viewing direction is directed towards the direction provided by the Miller indices.

The structure can still be rotated around the **z** axis without changing the orientation determined by **n**_*hkl*_. This remaining rotation is related to the choice of in-plane base vectors **v** and **w**. The software first chooses the two lattice vectors **v** and **w** in the plane denoted by (*hkl*), and thus perpendicular to **n**_*hkl*_, so that both vectors are as short and as orthogonal to each other as possible (see the blue cell in comparison with the yellow one in ***Figure 4***). Then the surface is rotated around the **z** axis until the longer of the two vectors **v** or **w** is aligned with the **x** axis. Since the lattice in the plane, i.e. the pattern of the points in ***Figure 4***, is dependent on the orientation of the plane, the choice of the base vectors differs for each choice of viewing directions **n**_*hkl*_.

Note that for odd combinations of large Miller indices, the UC vectors **v**, **w** and **u**—and with it the inclination UC—can get very large (many multiples of the size of the fundamental UC). ***Figure 4*** shows an example vector **v**, which is long compared to the size of the fundamental UC. This behavior directly translates to three dimensions.

Having the orientation fixed, the last step is to fix the translational degrees of freedom. This can be imagined as the oriented slice being a stencil, cutting a rectangular piece out of the infinite, periodic membrane structure at different locations. Moving a stencil along a periodic structure does not significantly change its content, given the stencil is at least the size of a inclination UC: the slice will in most cases contain one or multiple copies of the UC. As most biological samples will be ultra-thin slices, i.e. have a much larger base than thickness, this caveat is met in most cases in the **v** and **w** direction of the sample, i.e. its width and height, but not the **u** direction, the viewing direction. Thus, whereas translating the slice through the membrane structure in **v** and **w** direction does barely affect the projection, moving it along the **u** direction will impact the projection significantly, since different parts of the structure are contained in the slice.

SPIRE always centers the slice around the origin, however, allows translation along the normal vector **n**_*hkl*_. In case of a slice thickness smaller than the size of the inclination UC in the normal direction, this provides the ability to scan through different parts of the UC.

### Synthetic microscopy images as projections of the discretized membrane model

Based on the above virtual 3D model of the desired membrane structure, we now compute the planar projection and thus simulate the TEM image.

The planar projection of a membrane structure is a pixel image, thus an array of *n_x_* × *n_y_* pixels, each with a value indicating the brightness of the pixel. In the biological sample, a dark pixel in a TEM image indicates that much of the brightness of the incident beam has been lost due to a large amount of attenuation with matter, i.e. much electron-dense material is located along the path of the beam corresponding to that particular pixel.

This behavior is imitated in SPIRE: each voxel marked as being inside a membrane is assigned a numerical value of “1”, whereas all unmarked pixels, thus representing aqueous phases, are assigned a value of “0”. A value of “1” means that a membrane, therefore attenuating material, is located at that voxel. Adding all voxel values along a given direction yields the total amount of material interaction along that particular path. The values for each pixel at position (*i, j*) are computed by choosing a path along the viewing direction, thus normal to the *L*_*x*_*L*_*y*_ plane, where the path is located such that it intersects all voxels with identical and positions at *p*_*x*_ = *i* · *dx* and *p*_*y*_ = *j* · *dy*, as shown in ***Figure 2***. As a result, the final resolution of the projection is determined by the resolution of the voxel grid in the simulation box.

To avoid different brightnesses of projections due to varying numbers of voxels, all pixel values in the planar projection images are rescaled to an integer in the range between 0 and 255. This scaling allows for a linear or a logarithmic scale, which depending on the membrane structure may reveal more details in the projection.

### Additional structural properties

To provide further information about the projected 3D structure, the software computes the channel volume, the surface area of the membranes, and the percolation threshold of all channels (on demand). This section introduces, the underlying computational methods are introduced.

#### Channel volume

The computation of the channel volume is trivial, once the membrane structure is discretized. Each voxel occupies a volume of *V*_*υ*_ = *dx* · *dy* · *dz*, thus the total volume of a channel is just the number of all voxels associated to that channel multiplied with the volume of a single voxel. While there are discretization errors, these are small and decay quickly when voxel sizes are small.

#### Membrane surface area

The surface area of the membranes is computed using a triangulation of the membrane surface. The triangulation is computed by an algorithm called “Advancing front surface reconstruction”, implemented in the library CGAL (***Da and Cohen-Steiner, 2020***; ***The CGAL Project, 2020***), applied to the surface voxels of the membranes. The total surface area of the membrane is the sum of the areas of all triangles in the triangulation. As with the channel volumes computations, the quality of the approximation of the membrane surface by the triangulation, and thus the accuracy of the surface area, increases with an increasing number of voxels in the simulation box.

#### Percolation threshold

Percolation theory is an area in mathematics, statistical physics, and material science considering basic global connectivity properties of networks and graphs (***Stauffer and Aharony, 1992***). A network is said to percolate if a path through this network from a defined starting and end point can be found. Removing elements of such a percolating network can cause the connecting path to be cut and thus renders the network non-percolating.

This software uses percolation analysis to compute the percolation threshold of a channel, that is the maximum diameter of a body (e.g., a sphere or molecule) which can move freely through a channel of the entire structure without getting stuck at narrow passages (***Mickel et al., 2008***). This measurement is found by increasing the width of the membranes enclosing a channel step by step and checking if the channel is still percolating. The width at which the channel stops to percolate is called the percolation threshold, and denotes the most narrow diameter in the channel. A Hoshen-Kopelmann (***Hoshen and Kopelman, 1976***) algorithm is used to perform a cluster analysis of all voxels in a channel, i.e. all connected nodes of the network are grouped into a single object. The channel is percolating if only a single cluster is found during said analysis, that is each node (and thus voxel) in the channel can be reached from any other node in the same channel. A number of two or more clusters means that the channel has been separated.

### Identification of the structure of prolamellar bodies

#### Structure identification process

Here, we present the essential SPIRE features exemplified by matching TEM images of etioplast PLBs with software-generated projections. A video tutorial is provided, proposing an efficient workflow to recognize surface types with proper structural parameters visible on TEM images (http://chloroplast.pl/spire). The appearance of a 3D structure in a 2D projection depends mostly on i) the scale of a structure regarding its magnification, ii) the thickness of the visualized section, and iii) the orientation of a structure in the TEM sample. We discuss how to use these parameters for proper and efficient matching.

Since the size of the TEM sample is known, the first parameters which can easily be fixed to start the matching are the slice height, width and thickness. While the height and width only extend or reduce the size of the simulated sample, the thickness does influence the projection, as visualised in ***Figure 7***. Two examples of plastid cubic membranes varying in UC size and volume proportion of the two aqueous channels are provided: a gyroid membrane with a large UC and balanced channel volumes present in *Zygnema* sp. chloroplasts (***Zhan et al., 2017***) and a diamond membrane with a relatively small UC and imbalanced channel volumes found in bean *Phaseolus coccineus* etioplasts (***Kowalewska et al., 2016***). Drastic differences in the image characteristics point to the crucial role of the proper identification of scale and channel volume proportion of observed structures. All these properties can be easily calculated directly from the TEM micrographs using standard image analysis tools (e.g., ImageJ).

**Figure 7.**
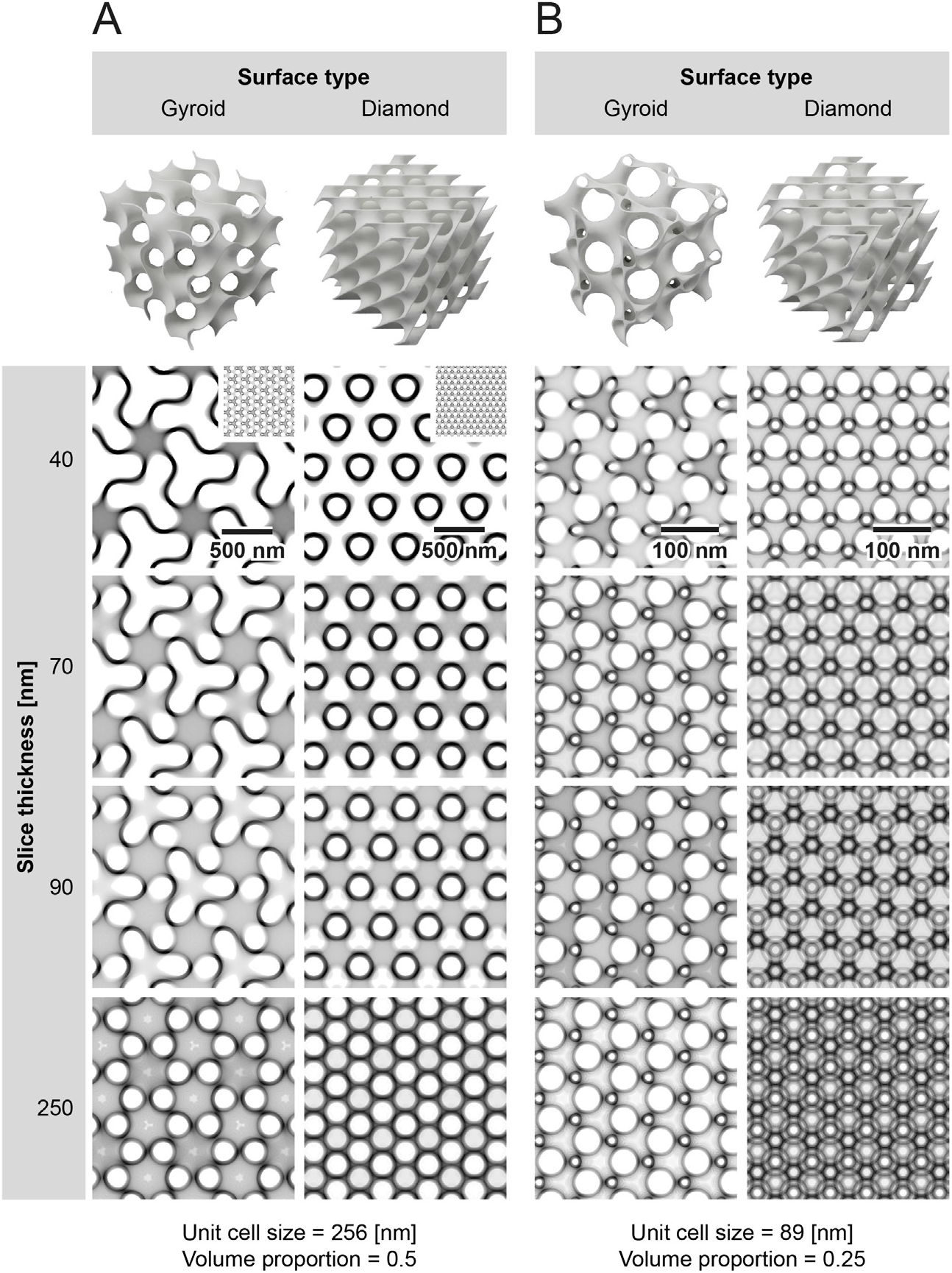
Diamond and gyroid type surfaces and computer simulation of Transmission Electron Microscopy (TEM) images of respective structures cut into sections of variable thickness (40–250 nm) The first row presents three-dimensional (3D) models of eight unit cells (UCs) of gyroid and diamond surfaces with balanced (**A**) and imbalanced (**B**) channel proportion. Computer simulations of TEM images are based on structural parameters (UC size and volume proportion – see bottom of the image) of cubic membranes recognized in (**A**) – gyroid of *Zygnema* sp. chloroplasts (*Zhan et al., 2017*) and (**B**) – diamond of bean *P. coccineus* etioplasts (*Kowalewska et al., 2016*). Note that for a better comparison, both surface types are simulated using the same structural parameters and are presented in (111) direction only. The figure shows how the slice thickness (subsequent rows) and structure scale (large (**A**) vs. small (**B**)) influence the pattern observed in computer simulations and, therefore, actual TEM images of such structures. Insets visible in the upper right corners of the second row of panel (**A**) present projections from panel (**B**) scaled equally; all parameters used to generate projections are listed in *Table 1*.

The next step is to make an initial (educated) guess for the structure type and orientation. A small gallery of all implemented surface types of different UC scales, volume proportions and orientations are presented in ***Figure 8***, which can be used to facilitate this step. Using the bulk creation functions in SPIRE, a user can also create their own, more refined and suitable galleries to provide better options for an initial guess. To create a gallery of projections and use it to match TEM images was originally proposed by (***Deng and Mieczkowski, 1998***), and this step in our software reimplements, automates, and improves this idea. The novelty and strength of our approach is the next step, in which the user can finely tune all parameters—with direct visual feedback—to further match the simulated projection to the TEM image.

**Figure 8.**
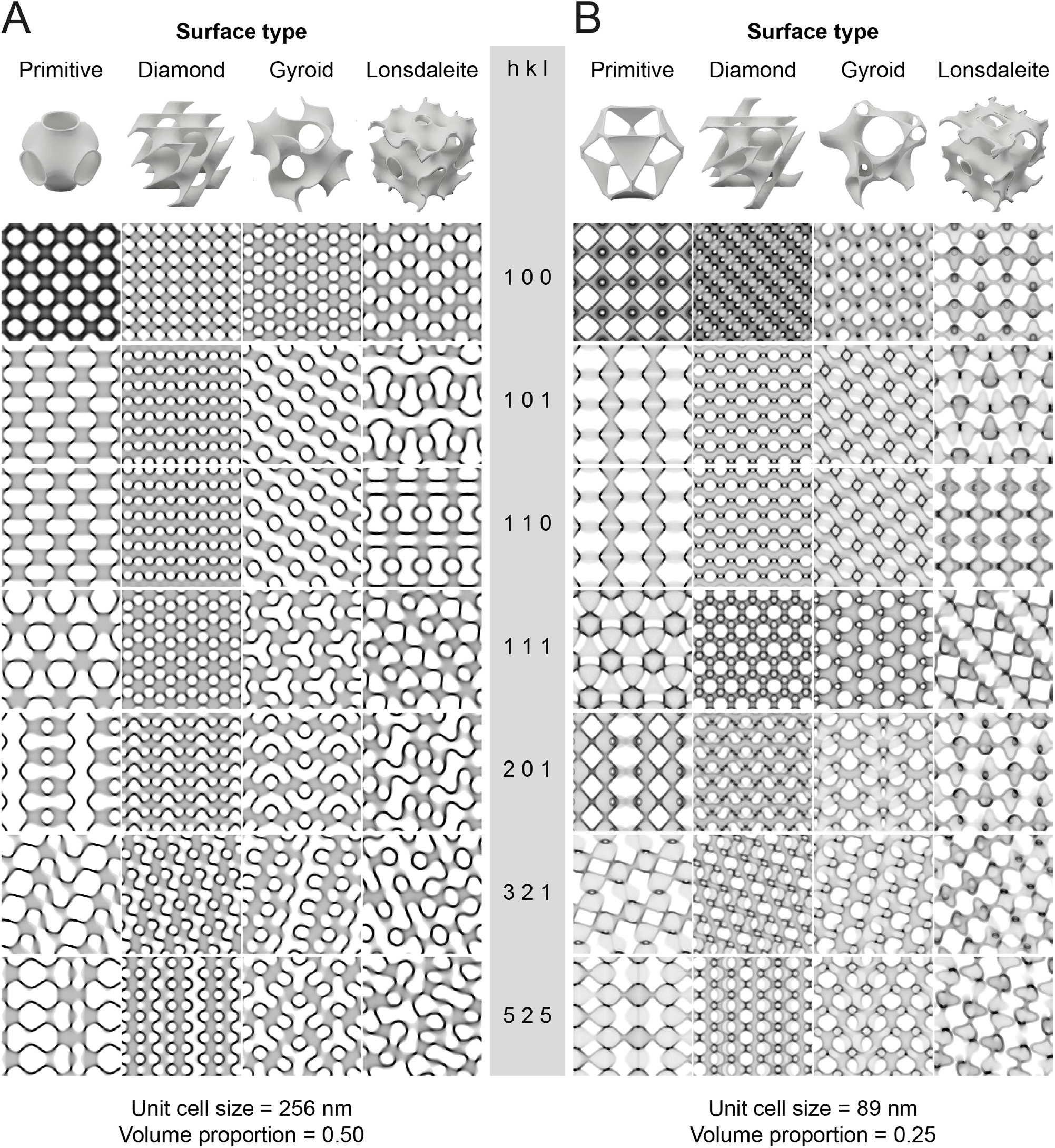
Gallery of selected (*hkl*) projections of four different surface types implemented in the software. The efficient process of surface matching is preceded by obtaining structural parameters such as unit cell (UC) size and volume proportion estimated directly from Transmission Electron Microscopy (TEM) images. The second step is completed by selecting a proper surface type and recognizing the structure’s orientation on the particular micrograph. For this purpose, a basic gallery showing variable (*hkl*) projections of different surface types, based on the idea ***provided by (Deng and Mieczkowski, 1998***), computed for balanced/imbalanced and large/small length scaled structures is a good starting point. Custom, more tailored galleries can be created by the user with the bulk creation function of the tool. Note that projections were computed to simulated TEM samples of 70 nm thickness for membrane structural parameters (UC size and volume proportion), same as in ***Figure 7*** on panels (**A**) and (**B**), respectively. three-dimensional (3D) models of periodic surfaces (first row) are presented for a single UC of all surface types; computed projections of TEM images are scaled to show the same number of UC despite the structure’s length scale; all parameters used to generate projections are listed in ***Table 1***.

If the slice thickness is not equal to the size of the inclination UC, i.e. it contains a fraction of an inclination UC (the UC in orientation), the projection differs depending on which parts of the inclination unit cell are contained in the sample. This effect is most impactful for very thin slices and declines the thicker the slice gets. In the software, the UC region is chosen by the slice position parameter. ***Figure 9*** shows three serial sections of the same PLB structure. Selected regions marked with different colors are matched with the following slices, taking into account the slice’s progressing position. The slight inaccuracies in the (*hkl*) values for the same regions in the subsequent slices are probably devoted to the sample warping during its visualization in TEM; note that neither the UC size nor the channel volume proportion was disturbed.

**Figure 9.**
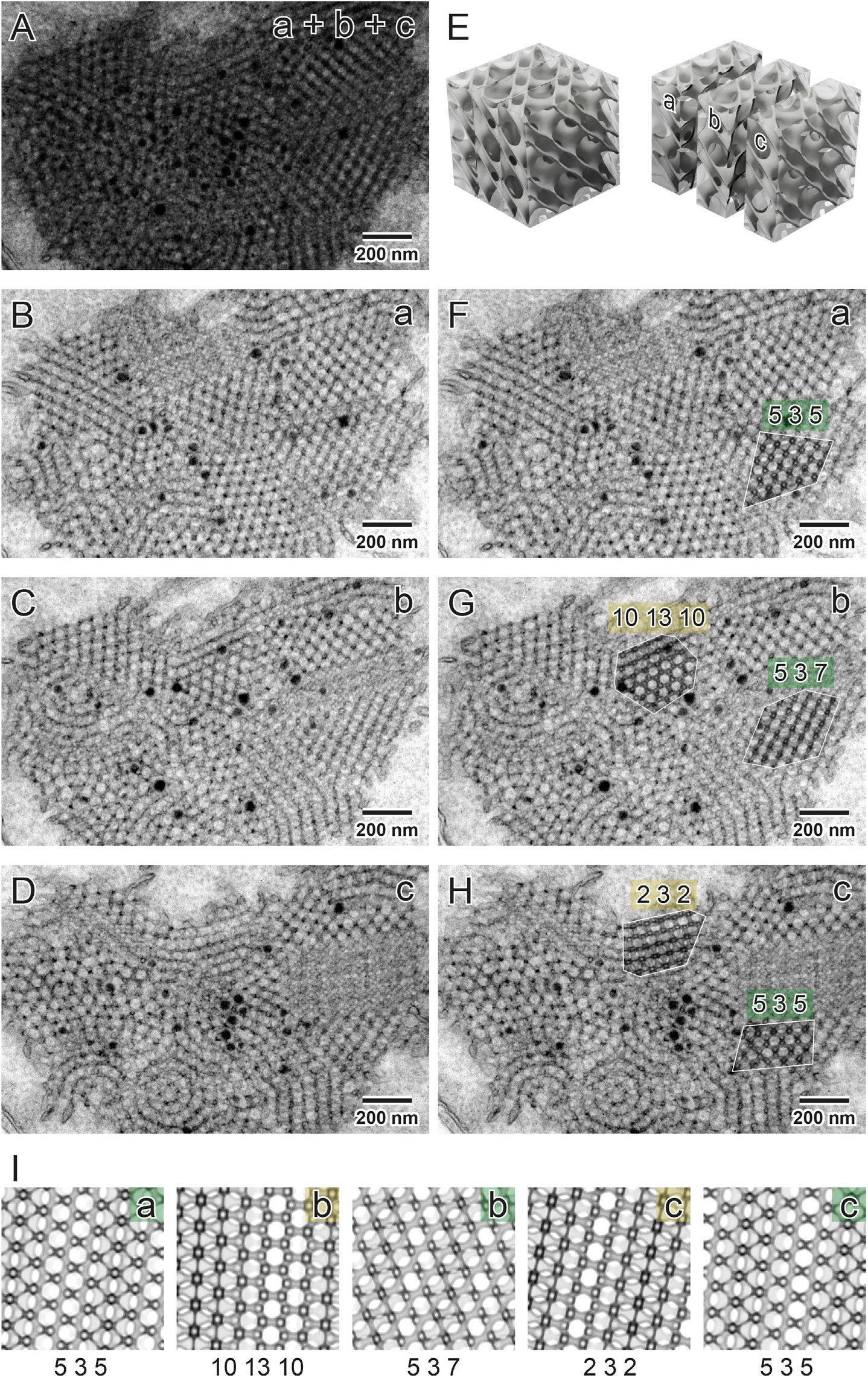
Surface type recognition in particular regions of oat *Avena sativa* prolamellar body (PLB) shown in serial Transmission Electron Microscopy (TEM) sections. Superimposed TEM images (**a+b+c**) showing PLB projections of a given thickness (70 nm) simulating a thick TEM specimen (210 nm) (**A**). Serial sectioning of a leaf sample enables visualization of subsequent regions of the PLB cubic structure (**B**-**D**); see an exemplary three-dimensional (3D) model of the cubic surface presenting the idea of parallel cutting of the specimen block (**E**). In principle, in selected regions (marked with yellow and green) of subsequent slices (**a**-**c**) matched projections should be the same (identical (*hkl*) values) but localized in the different depth of the structure. Such expected depth shift is observed in recognized projections (**I**) of a diamond surface type; however, matched projections are similar but not always identical in subsequent slices. Such an effect is probably due to the thin TEM sample’s warping during its visualization in the TEM chamber. Accuracy of projection matching is confirmed by superposition of computed projection and TEM image using multiply blend mode (F-H; regions marked with white border); all parameters used to generate projections are listed in *Table 1*

If possible, the identification of the structure should be confirmed by performing the matching procedure on several different regions of the sample with different orientations. These might be taken from different images or from a single image of a polycrystalline sample. PLBs often have a polycrystalline-like structure providing views of several projections of different orientations within one etioplast on a single micrograph; for details see ***Figure 10***.

**Figure 10.**
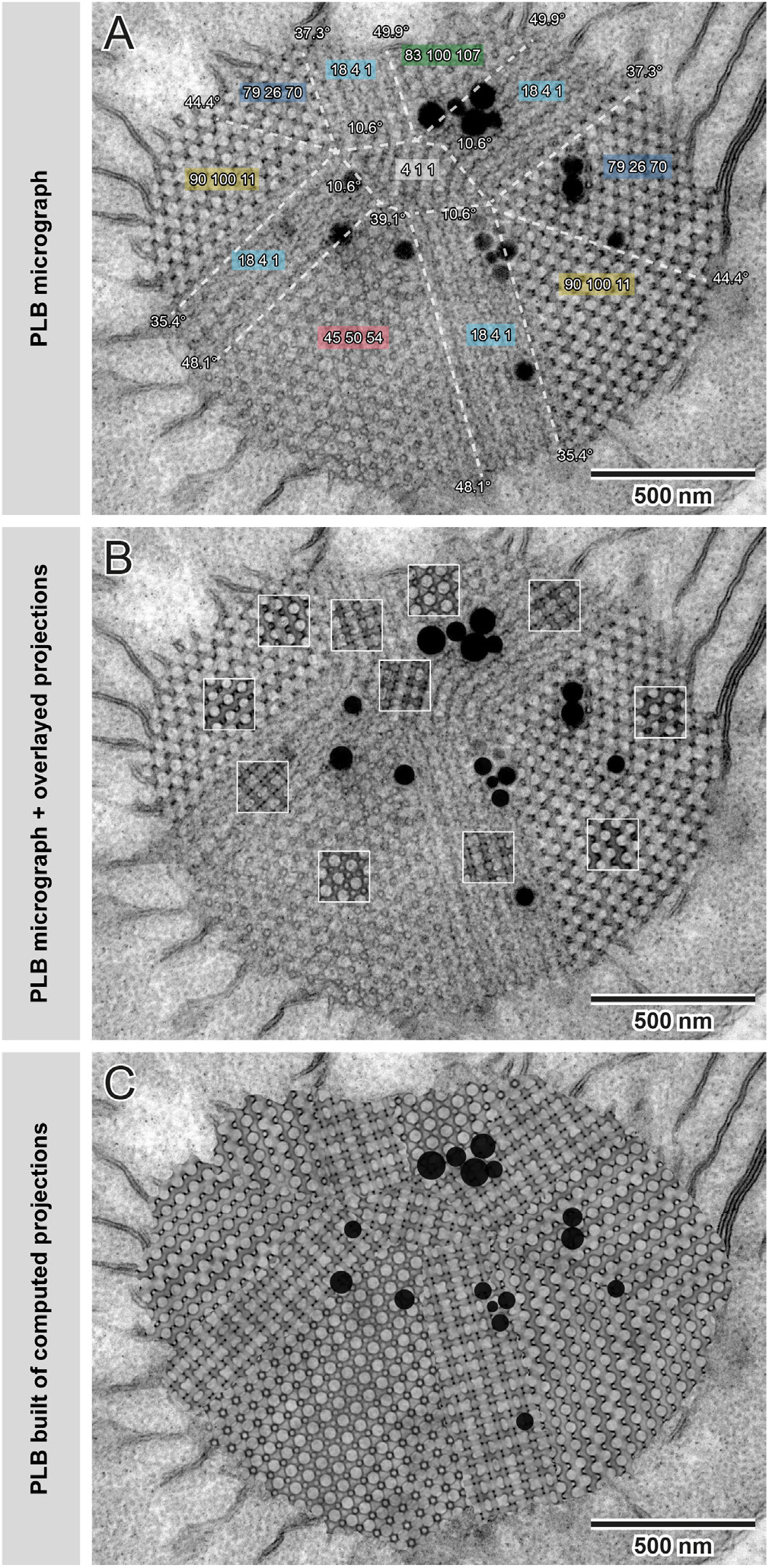
Construction of simulated oat *Avena sativa* prolamellar body (PLB) image built of computed projections. In many cases, PLBs and other naturally occurring cubic membranes are composed of several connected regions of bicontinuous surfaces at different orientations, forming “polycrystalline” arrangements (**A**). For a high confidence identification of the surface type, all different regions should be matched and analyzed. (**A**; same color indicates identical projections visible in PLB regions connected at different angles). To confirm the consistency of a match, the simulated projections can be superimposed on top of the Transmission Electron Microscopy (TEM) images (**B**; regions marked with white squares). To visualize the high accuracy of matches, we constructed an entirely simulated PLB built of particular computed projections (**C**) connected in the areas marked with white dashed lines visible on panel (**A**). Note that random noise was overlaid on computed projections to simulate the typical appearance of the TEM image; black dots added on the top of composed projections (**C**) indicate the position of plastoglobules visible on the TEM micrograph (**A**, **B**); all parameters used to generate projections are listed in *Table 1*.

The user can directly extract different structural features of identified surfaces from the measurement tab of SPIRE. Therefore, it is possible to automatically calculate variable 3D features of a recognized surface based on a 2D TEM image of the structure. Such information is particularly valuable from the biological point of view. Due to the lack of control over the direction of cubic membrane sectioning during sample preparation, recognition of surface type is based on different (*hkl*) projections. Calculations of channel diameters and UC sizes from 2D data are reliable only in cases of specific projections and in terms of primitive surface type also only in the exact depth of the slice. Therefore, the measurement functions of SPIRE, enabling calculations of the 3D features of the recognized structure, bring reliable information about, for example, the membrane area, the volume of the aqueous channels and the penetrability of the network by molecules of a given sizes; see percolation limit definition above.

#### Diamond as the predominant geometry in angiosperm PLBs

Using SPIRE and the described matching process, we identified the diamond surface type to be a dominating form of PLB cubic structures in the plethora of angiosperm species representing hypo- and epigeal germination as well as mono- and dicotyledonous plants (oat *Avena sativa*; ***Figure 9, Figure 10***, pea *Pisum sativum*, bean *Phaseolus coccineus*, cucumber *Cucumis sativus*, *Arabidopsis thaliana*, and maize *Zea mays*; ***Figure 11***). Although in all analyzed examples the PLBs matched the diamond surface, the UC size and volume proportion of both aqueous channels varied between 73.5–90.5 nm and 0.2–0.3, respectively (see ***Table 1***). In specific cases, the PLB can adopt a balanced diamond structure with a volume proportion reaching 0.5. Such a configuration has been so far identified in mutant plants with a disturbed composition of the PLB membranes only; e.g., *pif1 Arabidopsis thaliana* plants over-accumulating chlorophyll precursor (protochlorophyllide) (***Bykowski et al., 2020) (Figure 12***). Note that a significant increase in volume proportion alone, without changes in UC size, causes a rise in the membrane area packed in a given volume (***Figure 12***). Therefore, when the PLB size is maintained between different genotypes, such a balanced PLB network might store significantly larger amounts of membrane components, including enzymatic proteins and galactolipids crucial for efficient etioplast-chloroplast transition.

**Figure 11.**
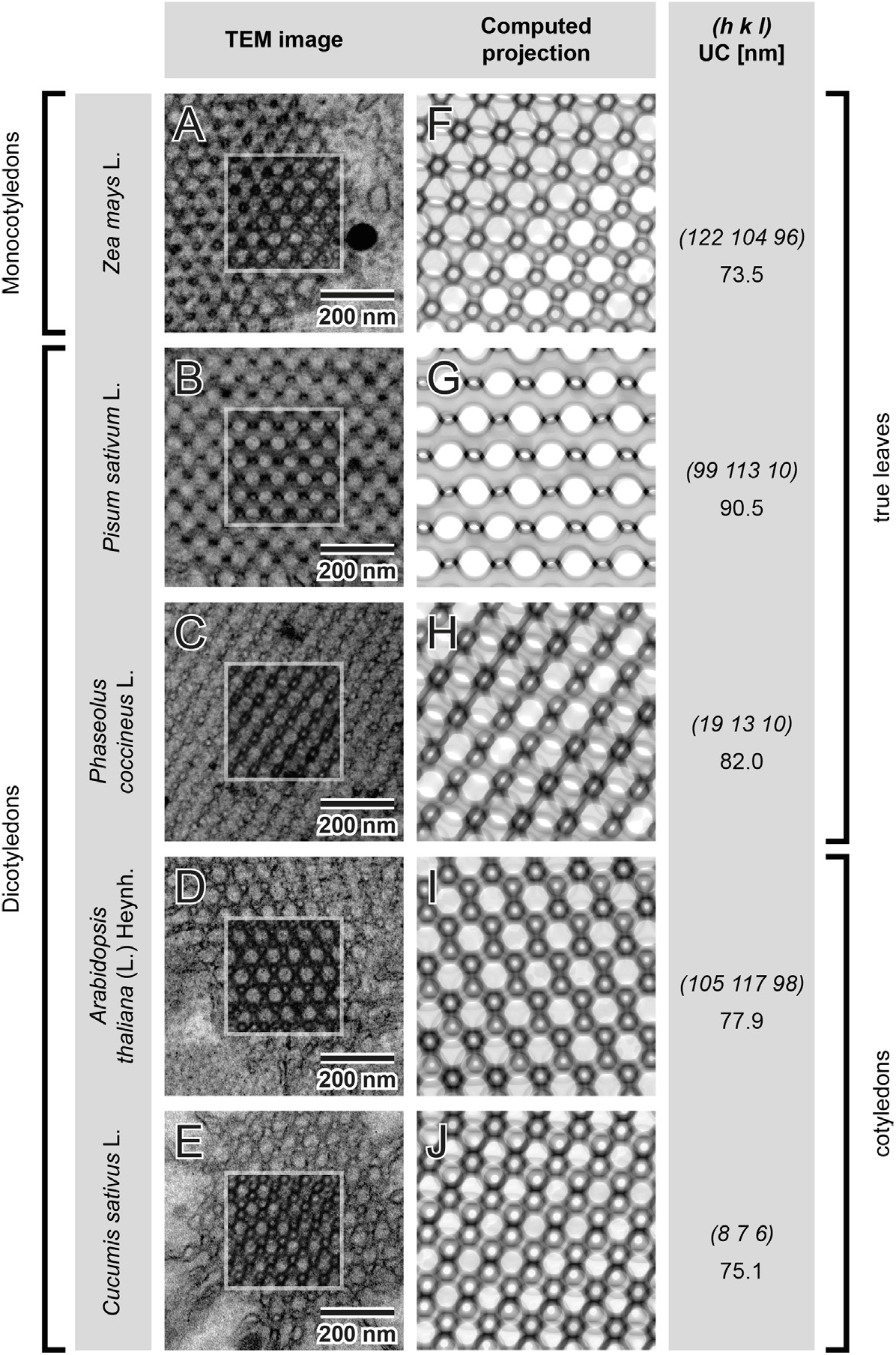
Diamond surface type—a dominating form of the prolamellar body (PLB) structure in etioplasts of angiosperms. Ultrastructure of the PLB in etiolated seedlings of several species from monocots (**A**) and dicots (**B**-**E**) exhibiting hypogeal (**A**-**C**) and epigeal (**D**-**E**) germination show diamond type of cubic structure. Regions marked with rectangles present superposition of computed projections and TEM images using multiply blend mode (**A**-**E**); all parameters used to generate projections are listed in *Table 1*.

**Figure 12.**
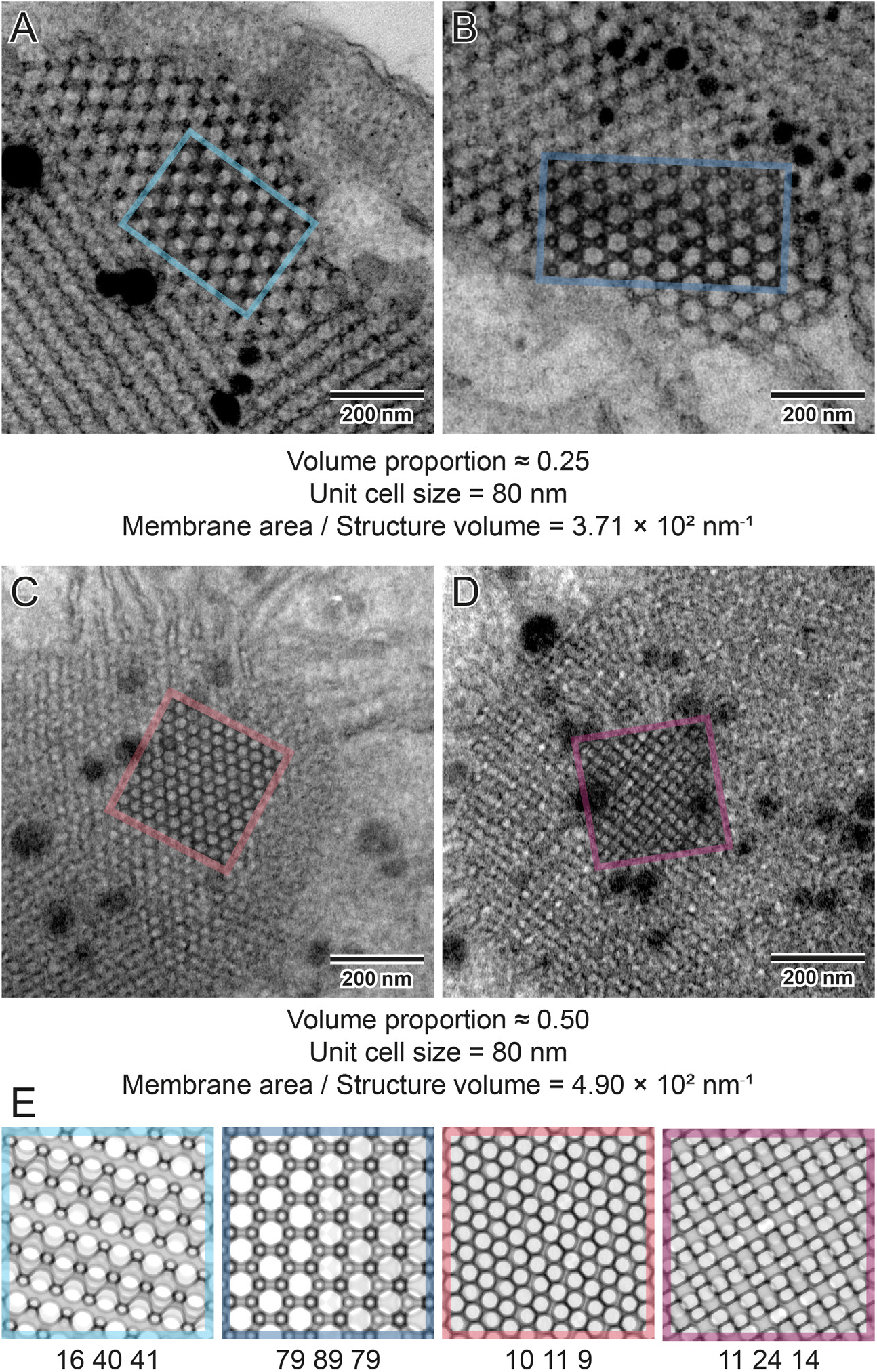
Imbalanced vs. balanced nature of prolamellar body (PLB) cubic membranes. PLBs are cubic membranes of diamond configuration characterized by an imbalanced structure in which two aqueous channels are geometrically different (**A**, **B**); regions marked with color rectangles show a superposition of computed projections and Transmission Electron Microscopy (TEM) images using multiply blend mode. Changes in the volume proportion resulting in balanced PLB structure without disturbances in the unit cell size are observed in plants over accumulating protochlorophyllide (*pif1* mutants of *Arabidopsis thaliana*) – a primary precursor pigment of etioplasts (**C**, **D**). Computed projections show how different are the observed patterns of networks that structurally differ only in the volume proportion ratio (**E**). This variability also influences the structure’s membrane packaging potential; balanced PLBs can accumulate more membrane components in the given volume; all parameters used to generate projections are listed in ***Table 1***

In ***Figure 13*** we present the exemplary utilization of the percolation threshold function.The obtained values indicate that the oat PLB structure enables a free flow of chloroplast ribosome particles through a larger aqueous network channel. Our percolation limit calculation stays in line with recent experimental electron cryo-tomography data, which confirmed the presence of fully assembled ribosomes at a stromal side of ruptured pea etioplast PLB (***Floris and Kühlbrandt, 2021***). However, it should be stressed that in both cases the size of the ribosome and the channel diameter are similar. Therefore, in PLBs of smaller UC size or varying volume proportion registered in different species, such mobility will be blocked. This suggests that ribosome presence in the PLB network may not be crucial for its proper functioning.

**Figure 13.**
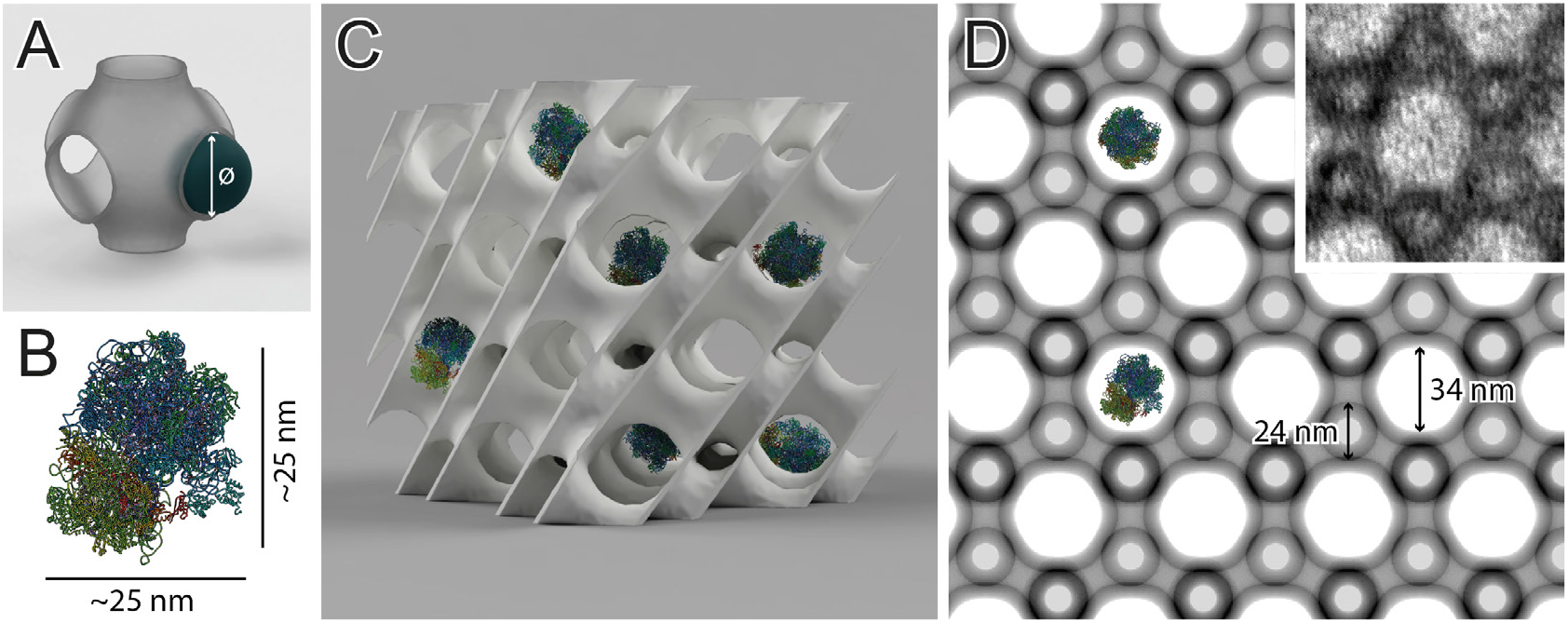
Penetrability of the three-dimensional prolamellar network – combining ultrastructural and molecular data. To estimate the maximum diameter of a sphere that can freely penetrate through a channel of the entire structure, the “percolation threshold” implemented in SPIRE can be computed (**A**). Here we used an example of plastid ribosome (**B**– PDB 5MMM, spinach *Spinacia oleracea* 70S chloroplast ribosome modeled using Chimera software) which size is comparable with diameters of the oat *Avena sativa* prolamellar body (PLB) channels. Using the percolation threshold function, it is possible to estimate whether a molecule whose spatial structure is already revealed could move freely through the channels of a particular cubic surface. The percolation threshold (31.57 nm) of a given network (unit cell size 80 nm, volume proportion 0.25), identified by a structure’s projection (**D**), can be calculated in SPIRE. The diamond network of oat PLB in (111) direction is presented with ribosomes in the same scale, region marked with white square presents superposition of computed projection and TEM image using multiply blend mode. Note that proper calculations are also provided for the surfaces recognized by projections in which the channel’s maximal diameter (40.11 nm) is not visible. In this example, ribosome size is smaller than the oat PLB’s percolation limit, which indicates that this molecule can move freely through the larger aqueous channel of the PLB network (stroma) of oat, hypothetically fulfilling its biological function directly inside the cubic structure; all parameters used to generate projection are listed in ***Table 1***.

## Discussion

In this work, we introduced a new software tool (SPIRE), which, based on “nodal surface” models, generates synthetic microscopy images of cubic membranes, bicontinuous phases and other structures. This tool enables the stereological identification of 3D structures based on their 2D projections, a key element in understanding structure-function relationships. We have demonstrated the basic concepts and workflow of SPIRE with a novel appliaction to one of the common examples of cubic membranes occurring in nature – etioplast prolamellar bodies. We revealed that PLB configurations resemble a diamond surface type and, despite earlier assumptions, is not based on a lonsdaleite (wurtzite) structure, at least for the plants analyzed here. Moreover, this work serves as a reference paper for the open-source SPIRE tool.

The development of the interdisciplinary field of naturally occurring cubic structures and their potential importance for designing lipid lyotropic crystals of remarkably variable functionalities relies on the availability of tools to analyze observed structures; tools which can be robustly used without a need for an in-depth understanding of the mathematical background. This field combines biology, physics, chemistry, and mathematics to introduce a structure-based approach in understanding biological processes and designing new functional materials.

At this stage, much of the experimental work of biologists does not contribute to the field and might even pass unnoticed due to the lack of a common recognition of cubic arrangements. This is particularly regrettable given that biological bicontinuous structures have achieved a property that remains largely elusive in synthetic cubic phases: structure sizes (lattice parameters) larger than 50 nm.

By expanding the pioneering work by ***Deng and Mieczkowski*** (***1998***), SPIRE fills a gap in the field of surface type identification of cubic membranes, which has only been partially covered by previous methods. The interactive matching process is manual, however, bulk creation capabilities of SPIRE enable a semi-automated approach.

While scattering methods have been used successfully to recognize bicontinuous phase surface types, these methods have a range of limitations in the context of biological samples. First, cubic membrane structures are, in many cases, restricted to certain regions of cells and, in a broader context, only particular tissues. In general, such sparse material does not generate sufficient signal in scattering techniques. Further, biological samples are characterized by considerable heterogeneity, which also excludes the potential of scattering methods application for the analyses of cubic structures of eukaryotic cells. The third limitation is directly related to the large scale of many already recognized cubic membranes; the size of UCs exceeds the range of dimensions recognizable via small-angle scattering techniques. Therefore, in terms of regular cellular networks based on TPMSs, the gold standard of structure recognition is set by electron microscopy studies. For this purpose, we developed SPIRE, which efficiently simulates microscopy images and enables direct input of data achieved from initial numerical analyses of the image completed outside of SPIRE—in any image analysis software which is available and familiar for the user.

In future work, we aim to improve and extend SPIRE capabilities. A prime target is to automate the matching process of the simulated projection with the actual TEM image. Here the toolbox of image analysis and classification can be employed. A very promising approach here are deep neural networks, specifically convolutional neural networks, which has been proven to efficiently classify images (***Krizhevsky et al., 2012***; ***Rawat and Wang, 2017***). Having several projections of a structure at different orientations availbale (as is the case for the samples presented here) could significantly increase the matching accuracy. SPIRE is ideally suited to generate training data sets of nearly arbitrary size for these purposes.

With SPIRE’s source code being designed to be usable as a library, as well as being available freely under an open source license, we also see the possibility to integrate the code into existing tools and software. This way, especially once the identification process has been fully automated, we hope to make the identification of structures from stereographic images available to a large and broad audience across many fields.

For now, simultaneously with this work, we provide a commonly available video tutorial (http://chloroplast.pl/spire) that provides a comprehensive guide for the efficient semi-automatic matching procedure and presents all basic functionalities of SPIRE, the measurement tab in particular.

An anticipated broad interest in periodic structure recognition in biological samples enabled by SPIRE will lead to the proper identification of many naturally occurring bicontinuous structures, including geometric measurements computed by the tool. Starting with and extending the tables and data of cubic membrane occurrence in biological systems provided by ***Almsherqi et al.*** (***2009***), ***Landh*** (***1996***) and further literature, this could be the start for a new repository, connecting geometric structures with their natural or synthetic occurrence and functions. By providing a rich interface and access options, such a database would open new opportunities for the meta-analyses of geometric arrangements occurrence in the contexts of their, e.g., composition, evolutionary background, developmental importance, and biological meaning, and thus pace the way for new insights on broader scales.

In most of the studies, the cubic membranes’ appearance is reported without further inter-pretation of the data nor their numerical analyses. SPIRE could help to re-evaluate and properly annotate numerous already published structures. SPIRE enables acquiring several spatial parameters of the network, which might also be interpreted in the context of other experimental data, e.g., percolation limit with the mobility of molecules of given sizes which presence in the aqueous environment of the network has been confirmed in biochemical studies.

SPIRE is a key tool to accelerate the dynamic field combining actual biological data, computer modeling, and, finally, obtaining synthetic periodic structures based on natural ones. Therefore, SPIRE has the potential to broaden our understanding of cellular cubic membranes, their biological role, and their relevance in designing nature-inspired artificial bicontinuous phases of comprehensive utilization.

## Materials and Methods

### Implementation Details

The tool (https://sourceforge.net/projects/spire-tool/) as well as the source code (https://github.com/tohain/SPIRE) and all dependencies are open source and thus freely and openly available. The software was entirely written in C++ providing an intuitive graphical user interface (GUI), shown in ***Figure 14***, implemented using the QT libraries (https://doc.qt.io/). Several libraries are used in this tool: integer math library (https://cs.uwaterloo.ca/~astorjoh/iml.html), openblas (https://www.openblas.net/), gnu multiprecision library (https://gmplib.org/) and gnu multiple precision floating point reliable library (https://www.mpfr.org/) are used to compute minimal UCs, zlib (https://www.zlib.net/) and libpng (http://www.libpng.org/pub/png/libpng.html) are used to output projections to the png image format, the computational geometry algorithms library (CGAL) (https://www.cgal.org/) is used to reconstruct surfaces to measure their area. Furthermore the algorithms from (***Felzenszwalb and Huttenlocher, 2012***) are implemented to compute the Euclidean distance map and the ***Hoshen and Kopelman*** (***1976***) algorithm is used to compute the percolation threshold.

A separation of the computational core code and the interface code allows the use as a library to incorporate into further projects. A simple command line interface for the creation of large batches of projections as an example is included.

The code is designed to allow an easy implementation of further surface types, given in the form of an implicit level-set equation.

### Plant Material and Growth Conditions

*Avena sativa* L., *Zea mays* L., *Pisum sativum* L., *Phaseolus coccineus* L., and *Cucumis sativus* L. dark-germinated seedlings were etiolated for one week in high closed glass containers on wet paper moistened with nutrient solution containing 3 mM Ca(NO_3_)_2_, 1.5 mM KNO_3_, 1.2 mM MgSO_4_, 1.1 mM KH_2_PO_4_, 0.1 mM C_10_H_12_N_2_O_8_FeNa, 5 μM CuSO_4_, 2 μM MnSO_4_ 5 H_2_O, 2 μM ZnSO_4_ 7 H_2_O, and 15 nM (NH_4_)_6_Mo_7_O_24_ 4 H_2_O, pH 6.0 to 6.5, RT. Seeds of *Arabidopsis thaliana* (L.) Heynh. mutant pif1 (N66041; ***Huq et al., 2004***) were obtained from The European Arabidopsis Stock Center. *A. thaliana* seeds were stratified in 4°C for 24 h, and 4 h illumination (120 μmol photons m^−2^ s^−1^ 23°C) was applied to induce germination. Seedlings were etiolated for 5 days in Petri dishes on Murashige and Skoog Basal Medium supplemented with Gamborg B5 vitamin mixture (M0231, Duchefa Biochemie) and 0.8% Phytagel™(P8169, Sigma-Aldrich) in 23°C. Leaf and cotyledon samples were collected under photomorphogenetically inactive dim green light.

### Transmission Electron Microscopy (TEM)

Leaf specimens were fixed in 2.5% glutaraldehyde in 0.05 M cacodylate buffer, pH 7.4 (prepared using 25% glutaraldehyde solution G5882, Sigma-Aldrich; sodium cacodylate trihydrate C0250, Sigma-Aldrich; pH adjusted with 0.1M HCl) for 2 h, washed, and postfixed in 2% OsO_4_ in 0.05 M cacodylate buffer, pH 7.4, at 4°C over-night. We applied glutaraldehyde as a preferable fixative for preserving cubic membrane structures (***Chong and Deng, 2012***). Samples were dehydrated in a graded series of acetone and embedded in epoxy resin (AGR1031 Agar 100 Resin Kit, Agar Scientific). The material was cut on a Leica UCT ultramicrotome into 70 nm sections. Samples were analyzed in JEM 1400 electron microscope (Jeol) equipped with Morada G2 CCD camera (EMSIS GmbH) in the Laboratory of Electron Microscopy, Nencki Institute of Experimental Biology of Polish Academy of Sciences, Warsaw, Poland. The PLB ultrastructural features were measured with the help of ImageJ software (***Abramoff et al., 2004***). The periodicity of 2D sections was calculated based on averaged values obtained from Fast Fourier Transform (FFT) of PLB cross-sections. PLB tubule diameters were measured manually based on each tubule’s outer limits in particular orientations of PLB cross-sections.

### *In silico* image generation and manipulation

Projections obtained using SPIRE were superimposed on TEM micrographs (where applicable) using multiply blend mode in Adobe Photoshop. In ***Figure 10***C image was obtained using Adobe Photoshop by deleting a portion of TEM micrograph and substituting it with projections generated using SPIRE. Random noise was added using Add Noise filter with Gaussian Distribution and Monochromatic settings on a uniform grey image (RGB 127 127 127), blurred using Gaussian blur filter and superimposed on the image using multiply blend mode. Meshes of 3D models were generated in Houdini using the level-set representation of the surfaces and rendered using Autodesk Fusion 360 software. 70S chloroplast ribosome from *Spinacia oleracea* L. was obtained from RCSB PDB (accession number 5MMM) and rendered using UCSF Chimera software (***Pettersen et al., 2004***).

## Author contribution

writing - original draft (TH, MB, GEST, ŁK), writing - editing/review (TH, MB, MS, ME, GEST, ŁK), designed the software (TH, MB), implemented the software (TH), designed computational solutions (TH, MS), provided resources (TEM images) (ŁK), supervision (ME, GEST, ŁK), methodology of biological samples (MB, ŁK), performed image analysis (MB), prepared figures (TH, MB, ŁK), conceptualization (TH, MB, GEST, ŁK)

## Acknowledgments

We are grateful to Prof. Yuru Deng for her outstanding work in understanding cubic (bicontinuous) membranes in cell organelles, and for her pioneering work in using nodal surface representations for the identification of cubic membranes in biology. We thank Prof. Deng for sharing her insight during our conversations at the University of Copenhagen in 2019, from which this project emerged. We are grateful to Tabea Rettelbach for support in creating the video tutorial. GEST acknowledges funding from the Australian Research Council under the Discovery Project scheme, through project DP200102593. ŁK acknowledges funding from the National Science Centre, Poland, under grant number 2019/35/D/NZ3/03904.

## Projection parameters

**Appendix 0 Table 1.**
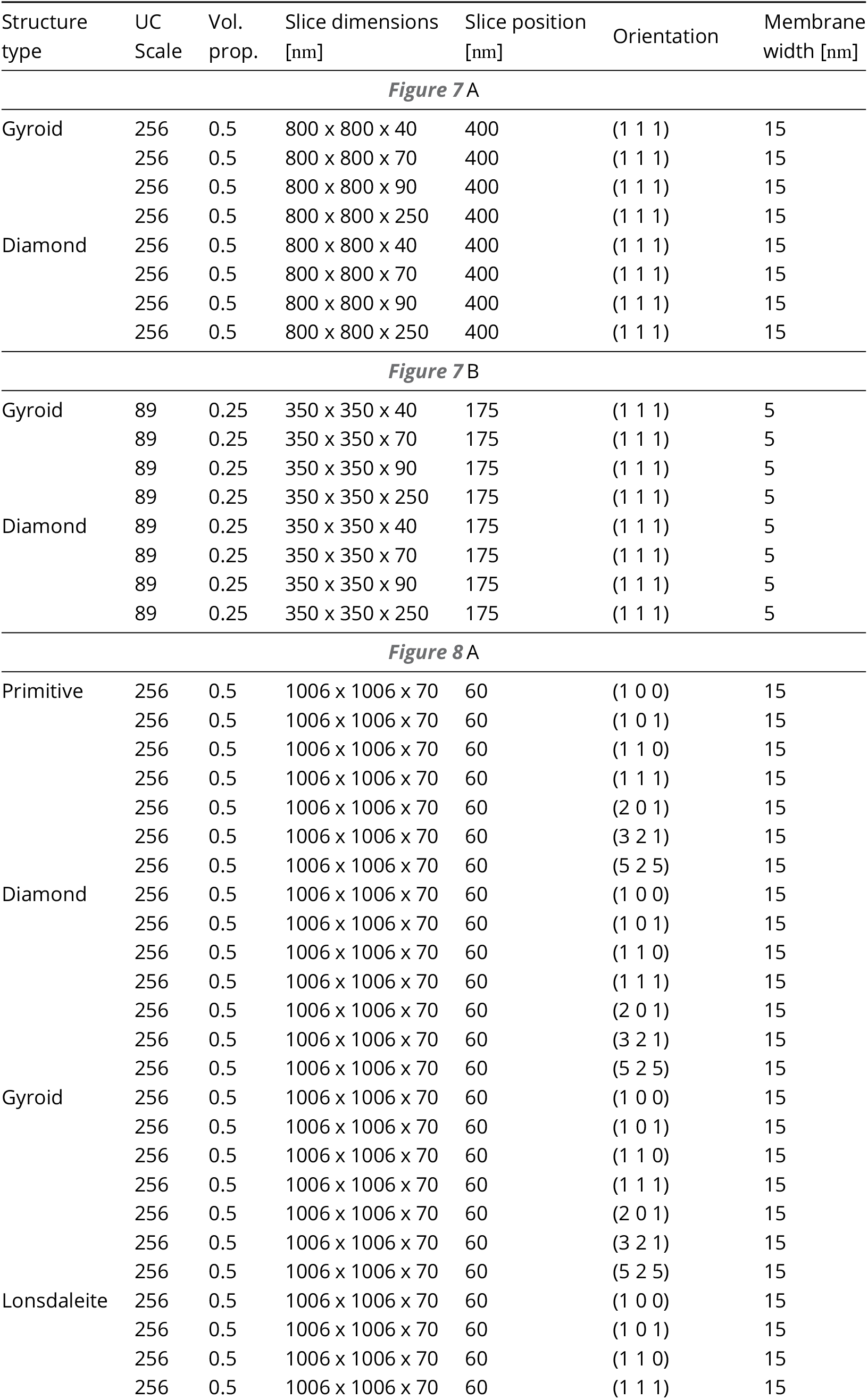
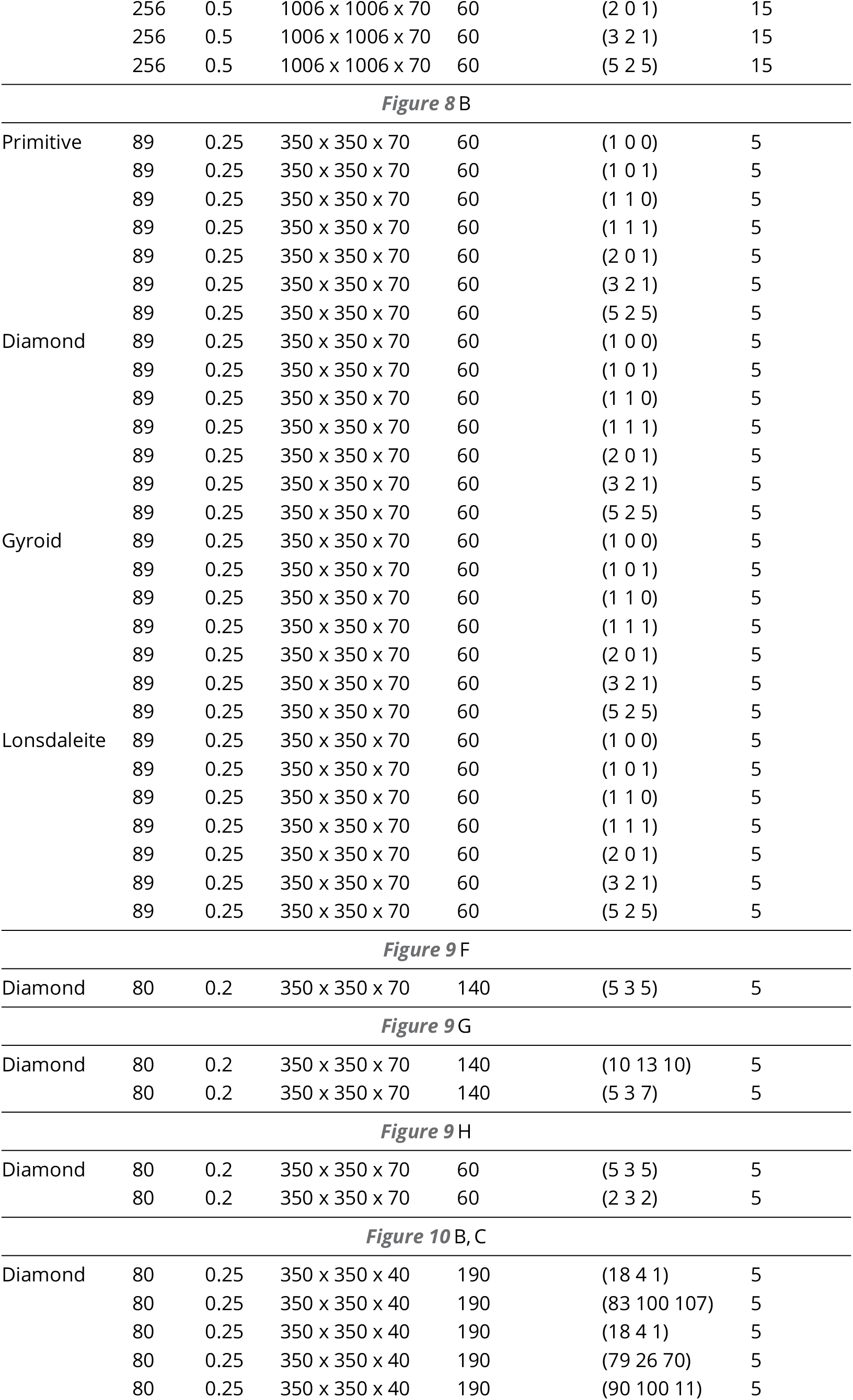
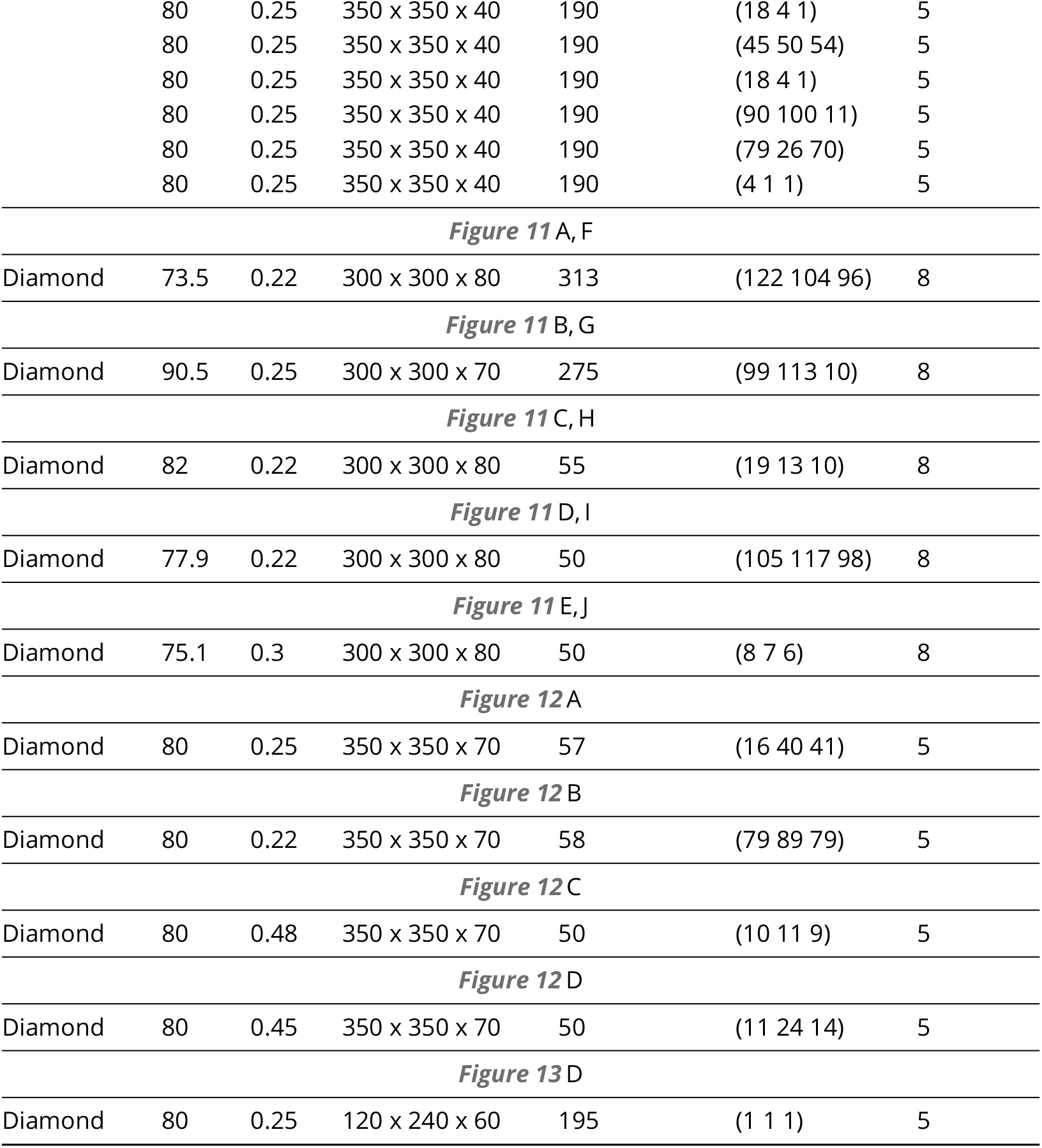
Parameters used to generate projections shown across the article including those matching actual TEM biological data.

**Appendix 0 Figure 14.**
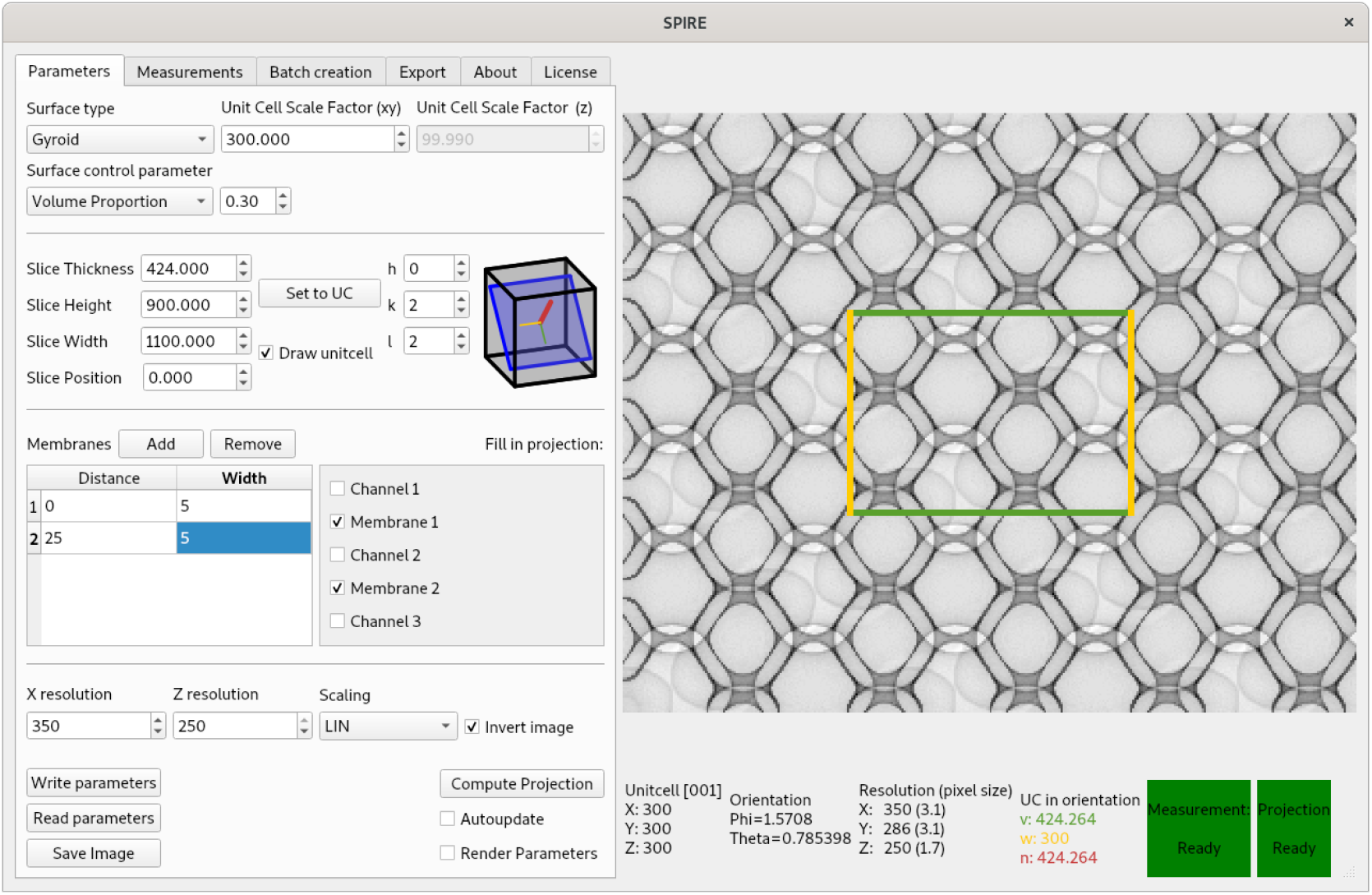
Screenshot of the Graphical User Interface (GUI) of SPIRE

## Fundamental unit cells of built-in surfaces

**Appendix 0 Figure 15.**
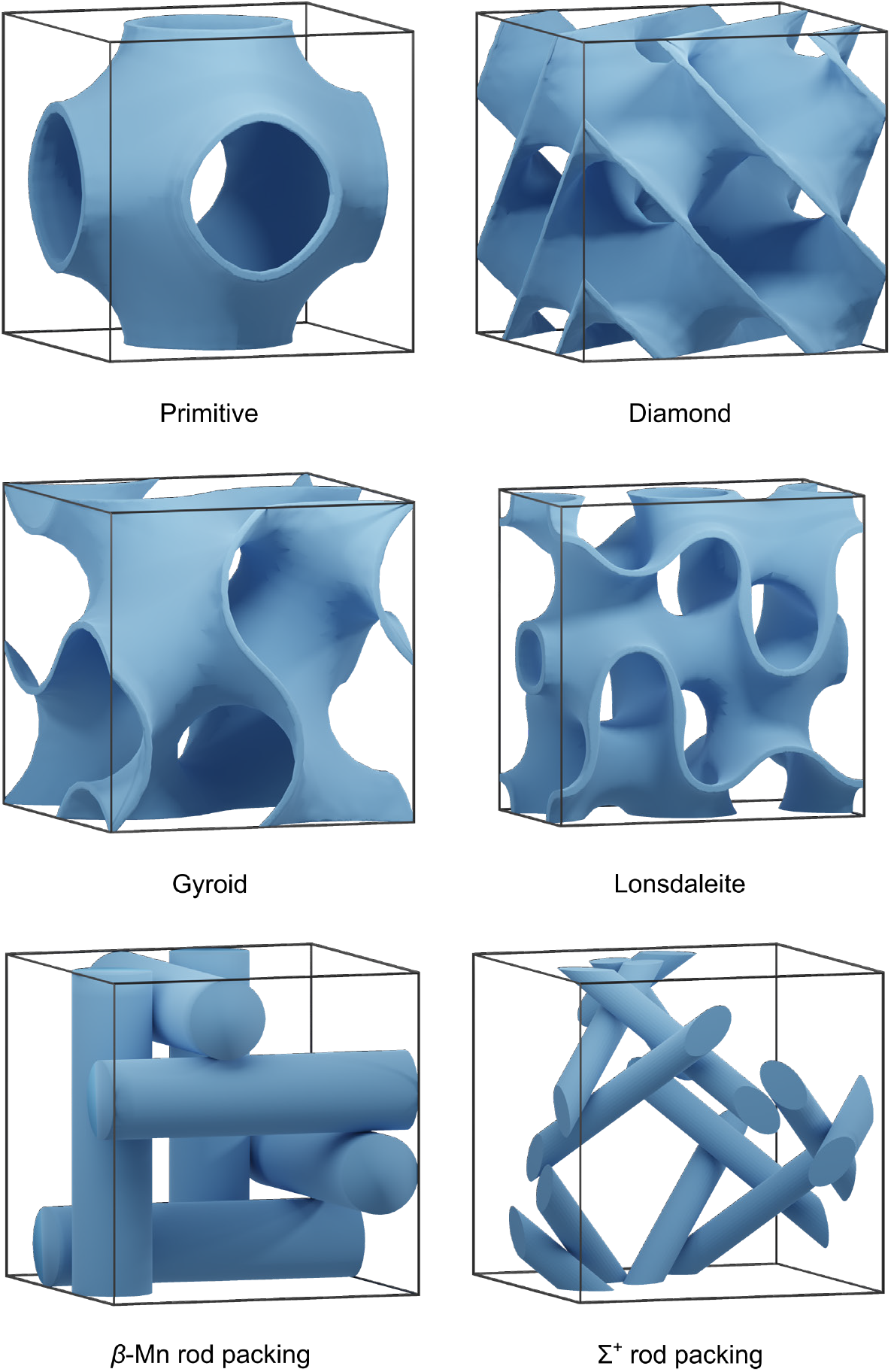
Renderings of the fundamental unit cells of the built-in structures

**Appendix 0 Table 2.** Choice of lattice vectors for fundamental unit cells

|  |  |
| --- | --- |
| Primitive | $a_1 = (1, 0, 0)^T, a_2 = (0, 1, 0)^T, a_3 = (0, 0, 1)^T$ |
| Diamond | $a_1 = (1, 0, 0)^T, a_2 = (0, 1, 0)^T, a_3 = (0, 0, 1)^T$ |
| Gyroid | $a_1 = (1, 0, 0)^T, a_2 = (0, 1, 0)^T, a_3 = (0, 0, 1)^T$ |
| Lonsdaleite | $a_1 = (1, 0, 0)^T, a_2 = (0, \sqrt{3}, 0)^T, a_3 = (0, 0, 1.732692)^T$ |
| Beta rod packing | $a_1 = (1, 0, 0)^T, a_2 = (0, 1, 0)^T, a_3 = (0, 0, 1)^T$ |
| Sigma rod packing | $a_1 = (1, 0, 0)^T, a_2 = (0, 1, 0)^T, a_3 = (0, 0, 1)^T$ |

**Appendix 0 Figure 16.**
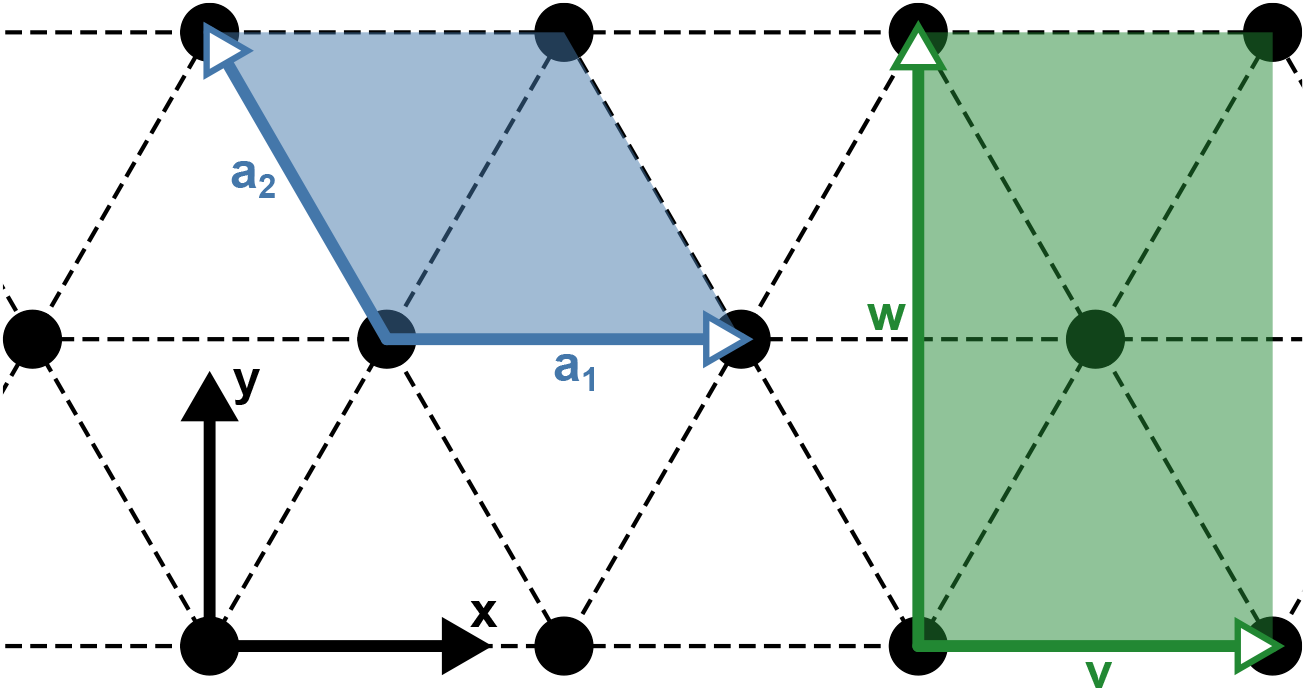
Choice of lattice vectors of the fundamental unit cell of the lonsdaleite surface. Shown is a top-down view of a hexagonal structure with the canonical choice of the unit cell (lattice vectors *a*_1_ and *a*_2_) and a rectangular unit cell (lattice vectors *v* and *w*). For convenience, we chose the rectangular unit cell over the canonical choice. The exact dimensions of the fundamental unit cell are provided in ***Table 2***

## Footnotes

1 A triangulated, minimal lonsdaleite surface was created using an input file by Ken Brakke for the surface evolver (***Brakke, 1992, 2008***). A voxelized version of the surface was then created by marking points on a rectangular grid as “1” on one side of the surface and “−1” on the other side. When interpolating, this grid has then the value “0” everywhere on the surface. This discrete, 3D test function was then approximated by a Fourier series *f* (*x*, *y*, *z*), using a Fast-Fourier-Transform (FFT). This Fourier series then has a root everywhere on the surface and for *f* (*x*, *y*, *z*) is thus a nodal representation of the surface. The same numerical protocol was used to compute nodal representations of two cubic rod packings, namely the *β*-Mn and Σ-^+^ rod packings. The original input was generated by placing cylinders of a given radius along the invariant axes of the rods, as described in ***O’Keeffe et al.*** (***2001***). Note that for convenience, in this implementation, instead of using the canonical choice of lattice vectors for a hexagonal symmetry, we use orthogonal lattice vectors with a rhomboidal symmetry. Please refer to the appendix Fundamental unit cells of built-in surfaces for the detailled information.

2 Whereas the surface *f* (*x*, *y*, *z*) = 0 will converge towards the true minimal surface by adding more terms to the series expansion, this is not the case for surfaces with *f* (*x*, *y*, *z*) = *c* where *c* ≠ 0. The latter, however are topologically equivalent (within a symmetric interval of *c* values) non-minimal, triply periodic surfaces.

3 There are cases, where structures grown onto substrates may exhibit preferred orientations, see ***Winter et al.*** (***2015***); ***Yoshioka et al.*** (***2014***)

